# Striatal Value Signals Track Progress Toward a Goal During Planning

**DOI:** 10.1101/2025.09.10.675322

**Authors:** Daniil Luzyanin, Arkady Konovalov

## Abstract

When making decisions, we often weigh immediate gains against long-term goals, and the value of the same choices can change over time. We used a novel task in which participants repeatedly chose between two cards, each with an immediate point value and an uncertain prospective value, with the latter reflecting the collection of card sets. Computational modeling showed that the prospective value captured how close participants were to completing a set. This goal gradient could be explained by both a simple progress-weighted heuristic and a forward-simulating planner. Using fMRI, we found that striatal and precuneus activity increased with the prospective value of the chosen card but not with its immediate point value. Activity in the dorsomedial and insular cortices instead increased with the value of the card given up and was highest when participants chose against the model-based value of the options. Activity in all regions of interest provides stronger support for the goal gradient rather than a full forward-planning model. These results refine the role of the valuation and cognitive control networks in goal-based decision-making.

## INTRODUCTION

Intelligent behavior often involves estimating the future value of our current choices, enabling us to anticipate outcomes and select actions that maximize prospective benefits^1–7^. When making decisions or planning, we often mentally simulate current and future actions and their consequences. This process has evolutionary advantages by helping us prepare for, or even influence, future events^8,9^. Understanding how the brain plans and estimates future states is important for many clinical applications. As an example, patients with prefrontal cortex damage struggle with multi-step problem-solving (for instance, the Tower of Hanoi task)^10^, and deficits in planning and future-oriented evaluation often contribute to impulsivity and maladaptive choices in disorders ranging from frontal dementia to substance abuse^11,12^.

One important open question is how the brain encodes current and future values of actions. The same choice options can have *immediate value* (e.g., intrinsic reward) and *future value*, which is the benefit the choice provides as a step toward a larger goal; we will call this value *prospective*. For example, taking a detour on a road trip may have a low immediate payoff (i.e., increased cognitive effort and longer travel time), but it can have high prospective value if it can help avoid traffic and save time later.

Converging evidence shows that the brain encodes value across a distributed network, with different areas possibly representing immediate or future outcomes. Much previous research on decision-making in this area has focused on how the brain values immediate rewards in simple, one-step choices, using time-discounting paradigms that do not involve explicit estimation of changing values but instead reflect an individual’s intertemporal preferences or patience^13,14^. These studies have identified a valuation network, including the ventromedial prefrontal cortex (vmPFC) and ventral striatum (particularly the nucleus accumbens), that encodes the subjective value of options at decision time^13,15–19^. Activity in these areas typically correlates with expected reward magnitude and/or probability, suggesting these regions may provide a common “currency” signal for comparing various options (items or actions)^17^.

This value network engages when choices involve trade-offs between immediate and delayed rewards. For example, it tracks the subjective value of future rewards discounted by a large delay, regardless of whether the reward is available now or will be received later^13^. At the same time, neuroimaging also suggests a degree of functional specialization in how the brain processes immediate versus future rewards. Some early neuroimaging studies reported a dissociation between choices involving delayed and immediate rewards, showing stronger limbic system activation for smaller immediate rewards and PFC activity linked to larger delayed rewards^14,20,21^. The valuation process in the PFC can also shape prospective choices and planning. Recent research indicates that damage to the vmPFC can impair planning depth^22^. Vice versa, prospective processes can also shape valuation. For example, imagining future events can reduce delay discounting by increasing coupling between the vmPFC/ACC (anterior cingulate cortex) and the medial temporal lobe^23^.

Another region often implicated in prospective decisions is the dorsal anterior cingulate cortex (dACC), which is involved in cognitive control and the selection of reward-guided actions^24,25^. The dACC is suggested to compute the expected value of control, estimating whether the potential future payoff of a difficult or costly action justifies the effort^24^. If the dACC integrates these signals, it can engage when the choice between immediate and future value becomes harder, with this trade-off often present during planning. For example, single neurons in the rodent and primate ACC have been shown to encode cost-benefit variables and future reward expectations during decision-making^25,26^. In humans, dACC activity has been linked to foraging decisions that require comparing the value of staying versus exploring for potentially better future rewards^27^. In general, the dACC and adjacent dorsomedial PFC (dmPFC) often show increased activation during tasks involving difficult, future-oriented choices, which could reflect the mental effort or conflict involved in choosing between short- and long-term goals^14,28^.

Two other key regions involved in valuation and planning are the striatum and the anterior insula. The ventral striatum (VS) often tracks the anticipated value of upcoming rewards^29,30^. The dorsal striatum (DS, particularly the caudate) can be engaged when actions are selected based on their consequences^31,32^ and has been shown to encode step-by-step values during forward planning^33^. In animals, striatal dopamine activity increases with the proximity and size of the reward as an animal approaches it^34–36^. The anterior insula (AI) is more associated with the salient and aversive aspects of choice options. For instance, activity in the AI increases with higher risk and anticipated loss^37,38^, with higher prices that lead people to forgo a purchase^39^, and with unfair rejected offers^40^. Insular activity is linked to interoceptive awareness^41^; together with the dACC, the insula is often described as a part of the salience network, which signals that behavior must be changed^42^. In meta-analyses of valuation, the anterior insula and dACC encode both gains and losses, suggesting a saliency rather than value signal^15^.

Planning in general is a complex behavior that involves computing both current and future values. Keramati et al.^43,44^ and Sezener et al.^45^ proposed that during planning the brain trades off the computational cost of simulating additional steps against the value gained by that foresight. These accounts suggest that planning depth increases when the prospective value of doing so is high, essentially, when the future rewards justify the effort. Studies of model-based reinforcement learning with multi-stage tasks have identified specific regions involved in forming prospective value (the caudate, lateral prefrontal cortex, and hippocampus) by correlating brain activity with computational model predictions^33,46–56^. First-stage choices in these tasks isolate prospective evaluation from value updating, even without the need to learn the task structure^57^. These studies linked striatal activity to model-free (learned through reward) and model-based (learned through the structure of the planning tree) prediction errors^46,57–59^ and OFC activity to a cognitive map of the task^60–63^.

Some of this work dissociated the neural systems involved in forward-looking evaluation from those involved in simple reinforcement learning. Using a task with several planning steps, Wunderlich et al. proposed distinct neural systems in forward planning versus choices resulting from overtrained habitual learning^33^. During planning, the anterior caudate encoded step-by-step future values, whereas the putamen tracked values learned through extensive training. Lee et al. identified a prefrontal arbitration mechanism that determines when to engage model-based planning^64^. They showed that the inferior lateral PFC and frontopolar cortex monitor the reliability of model-based vs. model-free predictions and allocate control accordingly.

While our understanding of prospective value encoding has improved, it remains incomplete. First, as discussed above, most human studies of prospective decision-making have focused either on purely immediate-reward choices (single-step preference tasks) or on paradigms in which future value is at least partly learned through trial and error (as in reinforcement learning tasks). These paradigms often require participants to learn a simple decision tree with a few stages, whereas in daily life the decision tree can often be too deep and complex to compute. Second, few experiments isolated neural signals of prospective value at the moment of choice, controlling for confounding factors such as reward prediction errors or ongoing trial-and-error learning.

Here, we used a novel task, computational modeling, and fMRI to address this gap. We developed a new card-collection task that dissociates immediate and future values within each decision, without requiring incremental learning. In this task, participants choose between two cards drawn from a deck; each card carries an immediate value (a point reward received immediately) and a prospective value that helps complete a “set” that yields a bonus at the end of the game. Participants are fully informed of the rules and payoff structure in advance. The task is stochastic, but the space of possible outcomes is so large that computing the decision tree exactly is impractical for a human decision-maker, although it can be approximated by simulation. Participants can still evaluate options prospectively, weighing whether a card’s immediate value is worth more or less than its contribution to a future bonus. Crucially, our design decorrelates the two value components across trials (r = 0.09 between the immediate and prospective value of the chosen card).

We investigated how participants compute forward-looking values in this paradigm. One possibility is that they use a forward model, approximately simulating how the rest of the game might unfold, and estimate how much each option increases their chances of completing a set. An alternative model is that participants rely on simpler heuristics, such as estimating whether a set of cards is still possible with each option, or how close they already are to completing a set, computing a “goal gradient” previously described in animal and consumer behavior^65,66^. Multi-stage planning can also be approximated by heuristics that truncate the decision tree^67,68^. We therefore compared several computational models, ranging from purely myopic choice to a normative planner that simulates the remainder of each game. This design let us ask three neuroimaging questions. First, do regions that encode subjective value represent the immediate or the prospective value of the presented options? Second, do regions implicated in conflict and control activate more when the two value components differ, or when a choice goes against the value of the option? Third, are neural value signals better described by a simpler progress heuristic or by a complex forward simulation?

**Figure 1.**
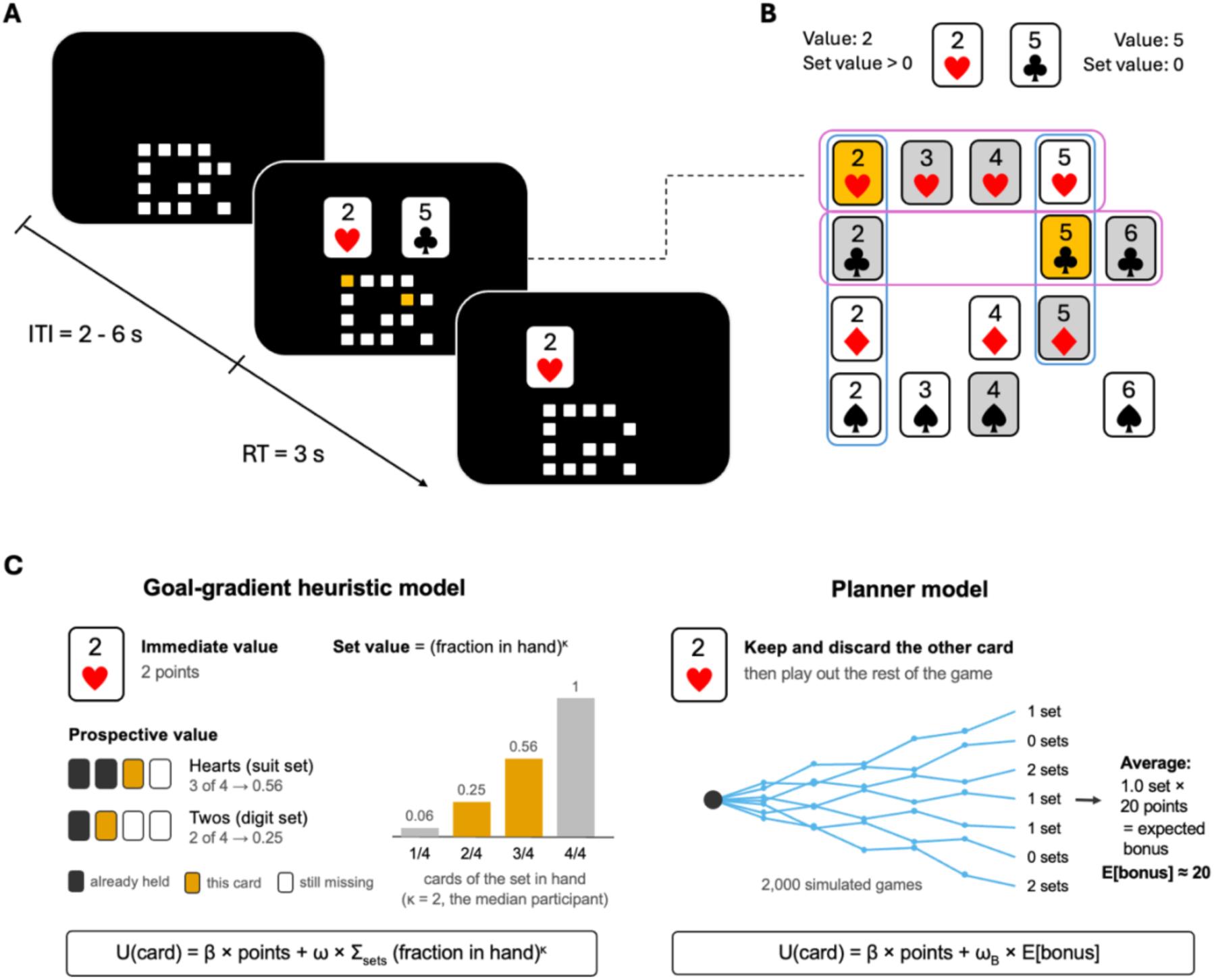
(A) Experimental task. In each game, the participant started with a deck of 20 cards, each representing four suits and five values (2 to 6), displayed in a grid at the lower part of the screen. All cards were greyed out at the start of the game. In each round, after a jittered inter-trial interval (ITI) of 2 to 6 seconds, two cards were randomly drawn from the deck, highlighted in the grid, and shown in full size above the grid. The participant chose one of the cards (left or right) using a button box. The response time was restricted to 3 seconds. The chosen card turned white, and the discarded card was removed from the grid. The game proceeded for 10 rounds, until all 20 cards had either been chosen or discarded. The display then showed the total points earned in this game. (B) The foundations of the decision problem. The computational models assume that the value of each card includes both its digit value (2 to 6) and its value as a part of a set. In the example shown, no more sets can be formed using the 5 of clubs, and two sets can be formed with the 2 of hearts. Thus, the 2 of hearts has a lower immediate value, but a higher prospective value. (C) How the two best-fitting models value a card (2 of hearts). The goal-gradient heuristic adds the value of each set the card can still help complete, which increases steeply with the fraction of the set in hand (shown for к = 2, the median participant). The planner model simulates the rest of the game 2,000 times and values the card by its expected set bonus.

## RESULTS

### Both immediate and prospective value contribute to decisions

We designed the task to distinguish between the immediate and prospective values of options (**Figure 1A**; see **Methods**). In the task, participants played a game using a deck of 20 cards that included four standard suits and five distinct digit values (2 to 6). During each trial, the computer drew two cards from the deck without replacement. Participants had to choose one card and discard the other. They immediately received the point value of the selected card, which we will refer to as “immediate value.” Depending on the luck of the draw, a participant could earn up to 52 points in a single game this way (four 6s, four 5s, two 4s).

They made ten decisions until the deck was exhausted. Then, they received an extra 20 points for each complete set of *at least four cards* sharing the same suit or point value (the set bonus was determined through pilot tests). Given the deck size, they could form up to three sets (for example, two suit sets and one set of digits), and the total possible set bonus could reach 60 points. Therefore, each card could have a “set value” (or, more generally, prospective value) if it could be part of a set. Participants often faced a trade-off between immediate and prospective value (**Figure 1B**), with the latter being crucial for successful planning and earning set bonuses. The game also featured long-term bonuses throughout the session (**Methods**), but these did not influence participants’ behavior. The weights on suit and digit sets did not change with the running difference between suit and digit sets collected (interaction with suit set weight: b = 0.10, z = 0.94, p = 0.35; with digit set weight: b = –0.06, z = –0.67, p = 0.50), and in model M5 suit and digit bonuses were weighted equally (Wilcoxon p = 0.20).

Always choosing the higher-value card is a suboptimal strategy in the task, and participants recognized that. Excluding ties (15.7% of trials, in which both cards had the same digit), participants chose the lower-point card on 25.7% of trials (SD = 13.3%), with considerable variability across individuals (8.8% to 54.4%). Participants also differed in task performance, earning between 1,789 and 3,384 points across the session (mean = 2,734). Participants who chose lower-value cards more often earned fewer points (r(27) = -0.48, CI = [-0.72, -0.13], p = 0.009), so overall, choosing higher values was beneficial despite the benefits of set collection. On average, participants earned 74.2 points per game. This was similar to the policy of always taking the higher card (74.3 points per game in simulation) and well below a simple policy that prioritizes feasible sets (83.7).

We found that participants’ choices relied on both the point values of the cards and their set values (**Figure 2A**). At every point difference, choices shifted toward the card that could still complete more sets. Here, a card’s set value is the number of sets (0, 1, or 2) it could still help complete in the current game. Participants recognized that and often chose a card with lower point value. Note that if this effect simply reflected uniform errors or low arousal, choosing lower value should be equally frequent across the set values. Instead, participants chose the lower-point card on 47.9% of conflict trials (where it had the higher set value), compared with 24.3% of neutral trials and 15.9% of congruent trials (conflict vs neutral: paired t(28) = 7.69, p < 0.001; neutral vs congruent: t(28) = 5.96, p < 0.001).

**Figure 2.**
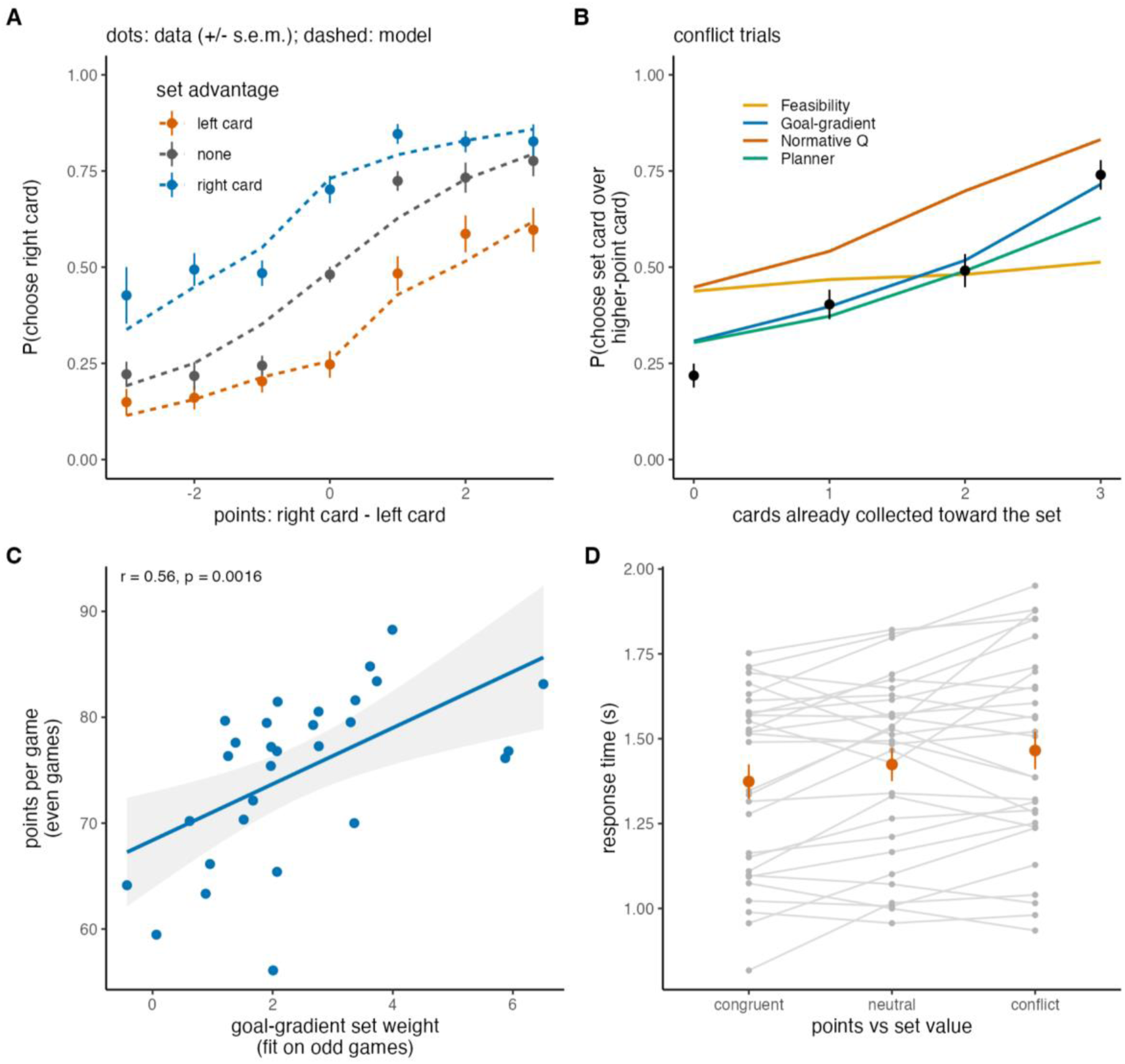
Behavior and computational models. (A) Probability of choosing the right card as a function of the point difference (right – left), split by which card had more feasible sets. Dots denote data (mean ± s.e.m. across participants); dashed lines denote goal-gradient model predictions. Individual participants are not shown for legibility. (B) On conflict trials, the probability of choosing the set card over the higher-point card as a function of how many cards of that set had already been collected (black dots denote data; lines show model predictions). (C) Goal-gradient set weight fitted on odd-numbered games against points per game in even-numbered games. (D) Response times on congruent, neutral, and conflict trials (gray dots are individual participants; orange dots are mean ± s.e.m.). N = 29.

### Prospective value follows a goal gradient that approximates planning

To explain participants’ decisions, we compared 8 computational models (see **Methods**). The goal-gradient heuristic (M3) fit the data best, and its summed BIC was 42 lower than that of the next best model, the planner (M4), and 49 to 1,144 lower than those of the other six models (**Supplementary Figure S1A; Supplementary Table S1**). This model was also the most likely model in random-effects Bayesian model selection (expected frequency 0.37, protected exceedance probability 0.83; planner 0.24 and 0.14; **Supplementary Figure S1B).** Model and parameter recovery showed that the models and their parameters could be reliably identified **(Supplementary Figures S1C and S2)** and the best model in leave-one-run-out cross-validation (cross-validated balanced accuracy 75.0 ± 10.8%, compared with 73.9% for M4 and 72.1% for M2; M3 vs M2: paired t(28) = 5.83, p < 0.001). The normative Q-value model (M7) fit the data the worst, suggesting that participants did not trade off immediate and prospective points one-for-one. In the planner model (M4), one point of expected set bonus was worth only 0.12 immediate points (median ω_B_/β; IQR 0.08-0.23; below 1 in 26 of 29 participants, Wilcoxon p < 0.001), suggesting that participants heavily discounted prospective value.

Model parameters predicted performance out of sample. We fitted the models to half of each participant’s games and related the parameters to the points earned in the other half. All weights predicted out-of-sample earnings. Goal-gradient set weight r = 0.63 and 0.56 (p < 0.001 and p = 0.002, for the two directions, **Figure 2C**), planner bonus weight r = 0.58 and 0.70, value weight r = 0.60 and 0.55 (all p < 0.03). Because all weights are also scaled by overall choice consistency, these correlations partly reflect less noisy choice in general, not only that participants relied on prospective value.

The two best-fitting models (goal-gradient heuristic and planner) mostly agreed on prospective value. Trial by trial, the goal-gradient set utility, and the planner’s expected bonus were highly correlated (r = 0.87). Most participants had a steep gradient (median к = 2; к ≥ 1.5 in 24 of 29 participants, set weight ω > 0: t(28) = 10.26, p < 0.001; **Supplementary Figure S3**). With к = 2, a set with three cards already collected is weighted sixteen times more than an untouched set. A qualitative test confirmed this (**Figure 2B**; see predictions of all eight models in **Supplementary Figure S4**). On conflict trials, participants gave up the higher-point card for the set card in 22% of cases when no card of the set had been collected, and in 40%, 49%, and 74% of cases with one, two, and three cards collected (mixed-effects logistic slope b = 0.82, z = 8.73, p < 0.001).

Simulating each participant’s full session with their own parameters, the goal-gradient model and the planner reproduced the observed number of sets per game (observed 1.51; simulated 1.52 and 1.53) as well as individual differences in sets (r = 0.92 and 0.95) and points (r = 0.94 and 0.96). The feasibility model under-collected sets (1.30 per game), and the normative model over-collected suit sets and chose the lower-point card too often (40% vs. 26%; **Supplementary Figure S5**). These results suggest that participants’ prospective valuation grows with progress toward a set and closely tracks the output of a forward-simulating planner, but participants estimate prospective rewards well below their true value.

Response times (RT), which were not used to fit the models, provided independent support for the modeling results. Participants were faster when the goal-gradient model predicted a larger absolute utility difference between the cards (log RT: b = –0.058 per utility unit, t(10.9) = –10.26, p < 0.001), as expected if difficulty reflects the computed values. Participants were slower on conflict trials (1,465 ms) than on neutral (1,424 ms; b = 0.039, t(27.3) = 2.40, p = 0.024) and congruent trials (1,374 ms; b = 0.060, t(26.2) = 2.98, p = 0.006; **Figure 2D**). Within conflict trials, choices of the set card were not slower than choices of the higher-point card (b = –0.028, t(30.2) = –0.97, p = 0.34). These results thus do not support the view that prospective choices require a separate, slower deliberative process and are consistent with a single integrated valuation, with conflict as the main factor for RT.

### Striatal and parietal activity tracks prospective, not immediate, value

We next asked how the two components of a card’s value are represented in the brain. GLM1 included, as simultaneous parametric modulators at the time of choice, the immediate and prospective utility of the chosen card from the goal-gradient model, as well as the utility of the unchosen card (see **Methods**). Prospective utility was associated with increased activity in the ventral striatum (right: x = 12, y = 5, z = –13; left nucleus accumbens: x = –12, y = 5, z = –10), the thalamus, the precuneus (x = –12, y = –67, z = 38), the left angular gyrus (x = –39, y = –52, z = 44), left dorsolateral prefrontal cortex (x = –45, y = 14, z = 44), bilateral orbitofrontal cortex (left: x = –33, y = 50, z = –13; right: x = 24, y = 53, z = –16), the supplementary motor area and the cerebellum (**Figure 3A**; **Supplementary Table S2**). No region showed a negative correlation with prospective utility.

**Figure 3.**
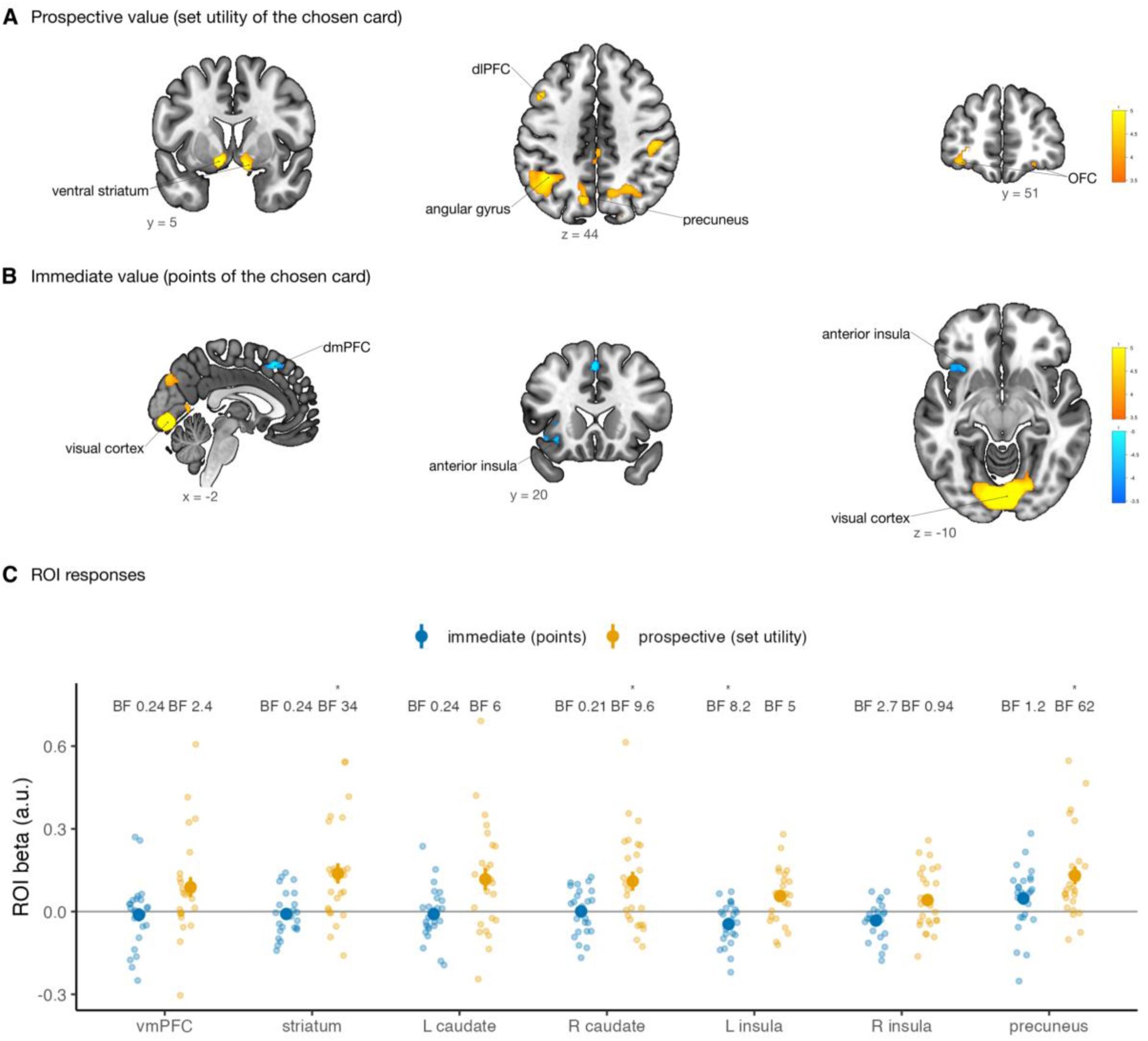
Prospective and immediate value of the chosen card (GLM1). (A) Regions where activity increased with the prospective (set) utility of the chosen card, including the ventral striatum (y = 5), the left angular gyrus, dorsolateral prefrontal cortex and precuneus (z = 44), and orbitofrontal cortex (OFC; y = 51). (B) Regions related to the immediate (point) utility of the chosen card: negative effects in the dorsomedial prefrontal cortex (x = –2) and in the left anterior insula (y = 20, z = –10), and positive effects in the visual cortex (x = –2, z = –10). Warm colors are positive effects; cool colors are negative effects. Maps are thresholded at cluster-level FWE-corrected p < 0.05 (cluster-forming p < 0.001; sign-flip permutation test). (C) ROI estimates for prospective (orange) and immediate (blue) utility. Small dots denote participants; large dots and bars denote mean ± s.e.m.; stars are Bonferroni-corrected p < 0.05 over seven ROIs; numbers are Bayes factors BF4o for immediate (left) and prospective (right) utility. N = 25.

The immediate utility of the chosen card was positively related to activity in the early visual cortex (x = 3, y = –85, z = –7; t = 11.3), likely indicating the larger number of suit symbols on higher-value cards. Striatal activity did not track immediate utility at all (**Figure 3C**). The immediate utility’s other effects were negative: in the dorsomedial prefrontal cortex (dmPFC; x = 0, y = 23, z = 50) and left anterior insula (x = –30, y = 20, z = –10) (**Figure 3B**).

Exogenous ROI analyses verified this dissociation (**Figure 3C**; all contrasts in **Supplementary Figure S6**). Prospective utility increased activity in the striatum (t(24) = 3.73, Bonferroni-corrected p = 0.007, BF4o = 34), the right caudate (t = 3.14, corrected p = 0.031, BF4o = 9.6) and the precuneus (t = 4.00, corrected p = 0.004, BF4o = 62), with moderate evidence in the left caudate (t = 2.91, corrected p = 0.053, BF4o = 6.0) and left insula (BF4o = 5.0), and weak evidence in the vmPFC (t = 2.43, corrected p = 0.16, BF4o = 2.4). Immediate utility had no positive effect in any ROI, and the Bayes factors favored no effect in the vmPFC, striatum, and both caudate nuclei (BF4o = 0.21-0.24). In the insula, immediate utility had a negative effect (left: t = –3.06, corrected p = 0.038, BF4o = 8.0; right: BF4o = 2.6). Prospective utility had a higher effect than immediate utility in the striatum and left insula (corrected p = 0.020 and 0.005), with weaker evidence in both caudate nuclei and the right insula (corrected p = 0.08-0.09). The difference between the utilities of the chosen and the unchosen card, the quantity most often reported as a chosen-value signal, was significant only in the precuneus (corrected p = 0.010, BF4o = 24); in the vmPFC, the evidence was inconclusive (t = 1.83, BF4o = 0.89). The weak vmPFC effects are unlikely to reflect signal loss as tSNR in this region was comparable to that in other ROIs and coverage was complete (**Methods; Supplementary Figure S7**).

### Dorsomedial prefrontal cortex and anterior insula respond when choices go against value

The negative relation between immediate value and dmPFC and insula activity could imply that these regions encode low immediate reward, or that they respond more strongly when the chosen card is worse than the alternative. GLM1 can distinguish between these two possibilities, as it also included the utility of the card that was given up. We found that activity in the dmPFC (x = 3, y = 32, z = 44), the right anterior insula (x = 33, y = 20, z = 2) and the right dorsolateral prefrontal cortex (x = 42, y = 23, z = 50) increased with the utility of the unchosen card (**Figure 4A**).

In GLM2, which included the signed decision value (utility of the chosen minus the unchosen card), the same regions showed negative effects. Activity was higher the more the chosen card’s utility was lower compared to the unchosen option (dmPFC: x = 0, y = 23, z = 50; right anterior insula: x = 30, y = 20, z = –10, t = –5.45; left anterior insula: x = –30, y = 26, z = –4; **Figure 4B**). The two maps overlapped in 112 voxels in the dmPFC and right anterior insula (Figure 4C). In the same model, activity in the dmPFC (x = 3, y = 41, z = 32) and bilateral anterior insula (right: x = 30, y = 20, z = –10; left: x = –30, y = 20, z = –7) increased with the unsigned decision value, the opposite of what a pure difficulty signal predicts. Together, these effects suggest that dmPFC and anterior insula activity were higher when participants chose the card with the lower utility, meaning they acted against the value the model assigned to the options. This pattern is open to several interpretations, which we consider in the Discussion section.

As model-derived utility includes set value, choosing against utility is not the same as choosing the lower-point card. GLM3 asked whether choosing the lower-point card engaged these regions differently on conflict trials, where card set value supported it, than on neutral trials. We found that it was not the case. None of the conflict contrasts produced effects in the dmPFC or insula, and in the insula ROIs the Bayes factors favored the null (BF4o = 0.24-0.42; **Supplementary Figure S6**). The only clusters that survived correction were in the left postcentral and supramarginal cortex and the right cerebellum (**Supplementary Figure S8**). As all responses were made with the right hand, these clusters, contralateral to the responding hand in the cortical areas and ipsilateral in the cerebellum, may reflect differences in motor execution between conditions rather than valuation.

**Figure 4.**
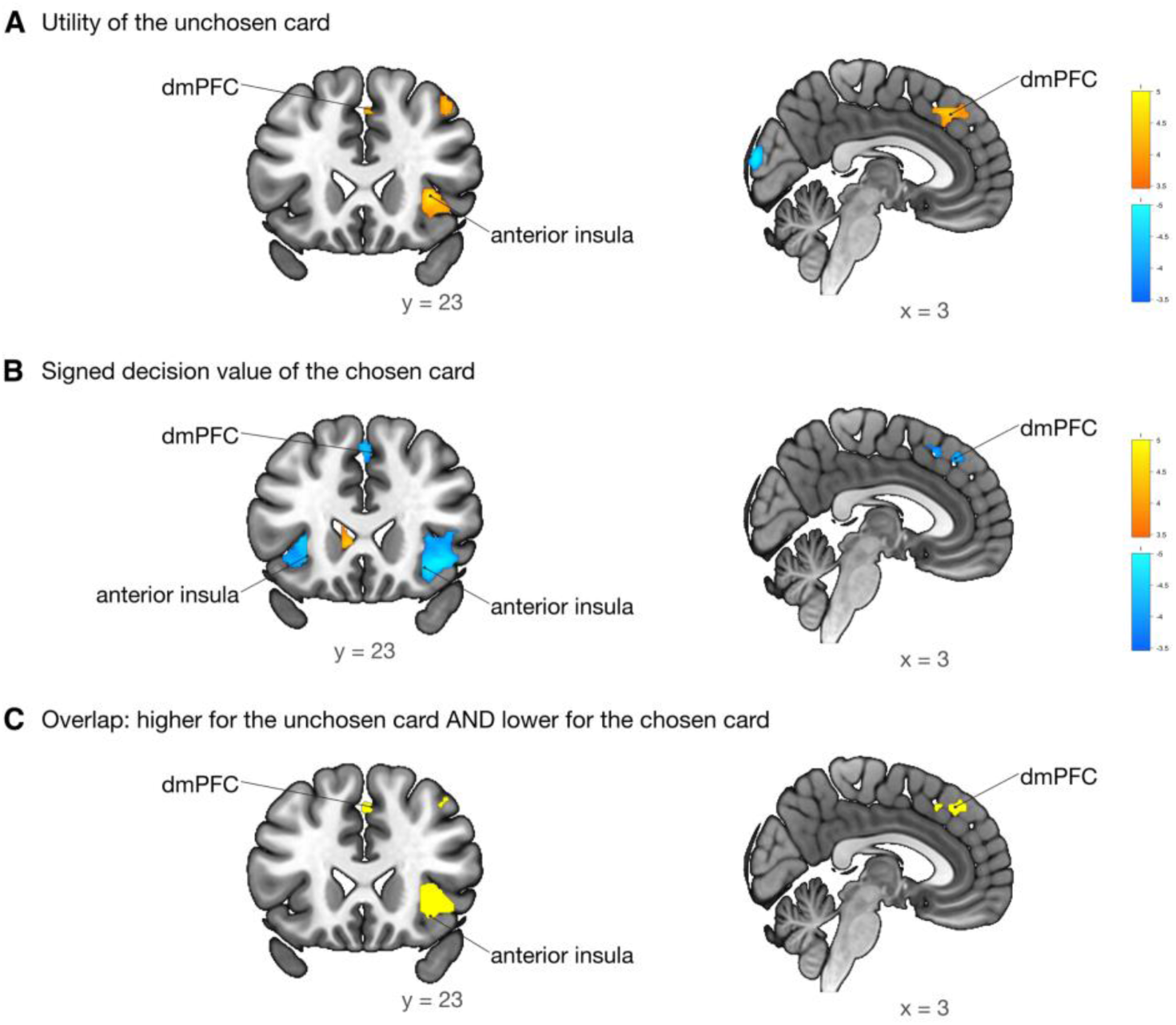
Choosing against value. (A) Activity related to the utility of the unchosen card (GLM1), positive in the dorsomedial prefrontal cortex and right anterior insula (y = 23, x = 3). (B) Activity related to the signed decision value, the utility of the chosen minus the unchosen card (GLM 2), negative in the same regions. (C) Voxels significant in both A (positive) and B (negative). Warm colors are positive effects; cool colors are negative effects; maps are thresholded at cluster-level FWE-corrected p < 0.05 (cluster-forming p < 0.001; sign-flip permutation test). N = 25.

### Neural model comparison supports the goal-gradient account of prospective value

We then asked whether neural prospective value is better described by the goal-gradient heuristic or by the planner model. GLMs 4 and 5 had the same structure and differed only in the prospective regressor: the goal-gradient set utility or the planner’s expected set bonus. Both identified overlapping networks in the ventral striatum and thalamus, lateral parietal cortex, and dorsolateral, frontopolar, and orbitofrontal cortex (**Figure 5A**). In the ROIs, both regressors were related to activity in the striatum, both caudate nuclei and the precuneus (all corrected p ≤ 0.03, BF4o = 10-137). Only the goal-gradient regressor was related to activity in the left insula (BF4o = 14 vs 0.34 for the planner) and showed weak evidence in the vmPFC (BF4o = 2.9 vs 0.29).

**Figure 5.**
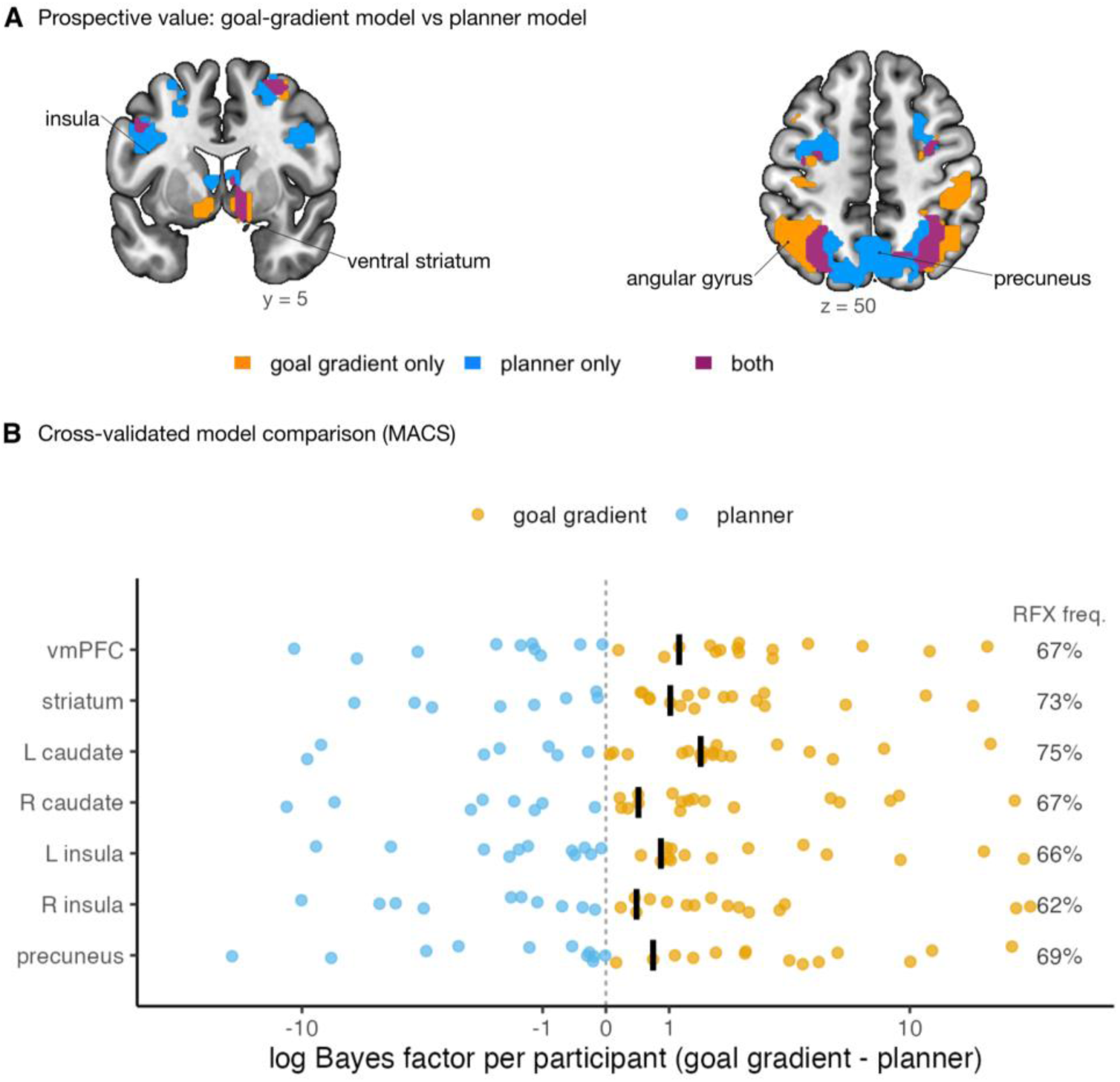
Goal-gradient and planner accounts of neural prospective value. (A) Regions where activity increased with prospective value computed by the goal-gradient model only (GLM 4, orange), by the planner only (GLM 5, blue), or by both (magenta); maps are thresholded at cluster-level FWE-corrected p < 0.05 (cluster-forming p < 0.001; sign-flip permutation test). (B) Cross-validated Bayesian model comparison of the two GLMs in each ROI (MACS). Dots show each participant’s cross-validated log Bayes factor for the goal-gradient versus the planner GLM (orange are participants with more evidence for the goal gradient model; blue are participants with more evidence for the planner; symmetric log scale); black bars denote medians; numbers on the right are random-effects expected frequency of the goal-gradient GLM. Dashed line denotes equal support. N = 25.

Cross-validated Bayesian model selection supported the goal-gradient GLM (**Figure 5B; Supplementary Table S3**). In all ROIs, more participants had stronger cross-validated evidence for the goal-gradient GLM (56-72%), its random-effects expected frequency was 62-75%, and the summed log Bayes factor was higher for the goal-gradient model in every ROI (+34 to +50). As the goal-gradient regressor used each participant’s estimated gradient steepness, whereas the planner regressor had no free parameter, part of this advantage may reflect that added flexibility.

## DISCUSSION

We found that participants’ choices followed a goal gradient, meaning a future opportunity mattered more the closer they were to achieving it. The best-fitting model was a heuristic that scaled set opportunities by progress toward the goal, but a planner that simulates the rest of the game and computes the expected bonus fit the data nearly as well. Participants deviated from optimality and valued expected bonus points at a fraction of their worth relative to immediate points. As a result, their earnings were no higher than those of a purely myopic strategy. Our results suggest participants’ behavior is consistent with a progress signal that approximates the output of forward simulation, consistent with accounts of planning as resource-rational approximation^67,68^.

### Valuation network and striatal encoding

We aimed to identify neural correlates of the immediate and future values of the same options using a new task that requires no reinforcement learning and in which exhaustive tree search is impractical. Our results largely align with canonical analyses of brain valuation networks^15,16^. However, they also reveal a dissociation between immediate-and prospective-value signals: activity in the striatum, caudate, and precuneus scaled with the prospective value of the chosen card, whereas Bayes factors favored the absence of an immediate-value signal in the same regions.

Our results match some model-based reinforcement learning findings, as studies on multi-step tasks have shown that striatal BOLD reflects not only model-free rewards but also model-based (future) value prediction^33,46^. Previous studies have shown that regions such as the vmPFC and ventral striatum encode a common “currency” of subjective reward^13,15^, typically responding positively to greater expected or received rewards. At the same time, work on intertemporal choice has highlighted a potential dual system: limbic-striatal areas respond to immediate gains, whereas lateral frontal/parietal regions are more involved regardless of immediacy^14^.

The striatal result connects to prior work on how dopamine and the striatum represent progress toward goals. Dopamine release in the ventral striatum ramps up as animals approach a distant reward^34–36^. Our prospective-value regressor was encoded in the ventral striatum at the time of choice. This result suggests that the striatal goal-gradient signal described in animals moving toward a reward may extend to abstract progress in a symbolic task. The caudate also tracked prospective value, consistent with its role in goal-directed action selection^31,32^ and with caudate encoding of forward-planned values^33^.

Neither the ventral striatum nor the caudate tracked the immediate points of the chosen card, and vmPFC showed only weak evidence for prospective value (BF4o = 2.4-2.9), no immediate-value signal, and inconclusive evidence for the difference between the chosen and unchosen card. This is unlikely to reflect potential signal loss, as tSNR in the vmPFC was similar to that in the other regions (**Supplementary Figure S7**). The vmPFC may therefore contribute less to the valuation of set progress than the striatum, at least at the time of choice. One explanation for the absence of the chosen value signal is that the vmPFC may evaluate or consolidate outcomes rather than participate in the planning process itself. Future studies could aim to dissociate the roles of this area using a more elaborate design.

### The dmPFC and insula: choosing against value

In our task, dmPFC and anterior insula activity decreased with the immediate value of the chosen card, increased with the value of the card that was given up, and was higher when participants chose the card with the lower model-derived utility. These regions therefore do not simply represent low reward. Similar dmPFC/insula responses have been reported in human fMRI when participants forgo a sure reward to acquire information or pursue future goals^24,69^.

Other interpretations are still possible. Choices against model-derived value are, by definition, those that the model predicts least well. The dmPFC and insula could therefore respond to response conflict, to monitoring a choice that departs from the prevailing policy, as in conflict- and error-related dmPFC activity^24^, or to exploration, or to occasional errors, and part of the effect could reflect trials on which the model’s predictions are wrong.

The dorsomedial prefrontal cortex (dmPFC) and anterior cingulate cortex (ACC) are often described as part of the cognitive control or salience networks. They are often recruited when difficult trade-offs or cognitive conflicts arise^24,25^. Critically, these regions are suggested to compute the expected value of control, weighing whether it is worth exerting effort now to secure future payoff^24,27^. Typically, the dmPFC/dACC activity increases when an immediate reward is forgone for a larger, delayed, or uncertain outcome. Likewise, the anterior insula co-activates with the ACC when individuals face risk, loss, or negative affective states, possibly signaling aversive aspects of a decision ^42,70^. The anterior insula result fits its proposed role in salience and in the aversive side of choice^15,30,38–40^. In our task, the insula may therefore register what is given up when people act against the more valuable option, a signal that could help to adjust later choices. The left insula also showed moderate evidence for a positive relation with prospective value (BF4o = 5.0), so the insula may carry both the value of progress and the cost of the choice; separating the two would require designs in which forgone value and prospective value vary independently.

### Implications for planning and model-based decision-making

Our findings have implications for theories of planning and value-based choice. First, the dissociation between striatal and dmPFC/insula signals indicates that the brain may use separate valuation channels for different temporal or prospective components of a choice. The striatum appears to carry the prospective component of value, consistent with forward-looking evaluation, which our modeling suggests may be implemented as a goal-gradient approximation rather than explicit search^4,46^. At the same time, the dmPFC and insula respond when the chosen option falls short of the alternative, which may help to monitor or override the prevailing value signal. This complementary pattern aligns with the idea of parallel decision systems^71^ but suggests a potentially more integrated implementation. Rather than describing one area that controls model-free computations and another that evaluates model-based plans, our results could support a model in which the canonical reward areas (striatum/vmPFC) encode the prospective value of each option, while control regions modulate behavior according to expected gains^24^. Our approach, however, differs from the common two-step and related paradigms^33,46^, which infer model-based activity through choices and prediction errors.

Although dopamine is typically associated with immediate reward prediction errors ^72^, some work^33,73^ shows that increasing dopamine can enhance model-based control. Potentially, the striatum is not limited to habitual “learned” values but can flexibly incorporate planned outcomes^53,63^. This idea suggests that “model-free” circuits may, in fact, have access to predictive representations (the successor representations or learned maps) that enable them to propagate future value without exhaustive tree search.

## METHODS

### Preregistration

We preregistered the study, including the task code, study design, and main hypotheses, on OSF (https://osf.io/f75gx). We chose the target sample size based on a previous study from our lab^52^, aiming to detect a large correlation between individual behavioral and neural measures (r = 0.5, a = 0.05, 80% power), resulting in a target sample of 29 participants. We did not preregister the computational models. As a main deviation from the preregistration, we included both value regressors in the same GLM for a more conservative analysis.

### Participants

We recruited 29 adult participants (18 female; average age 24.6 years, ranging from 18 to 35) from the Center for Human Brain Health participant pool at the University of Birmingham. All participants were free of neurological or psychiatric disease, right-handed, and had normal or corrected-to-normal vision. Participants received a fixed fee of £30 plus a bonus of up to £10 based on their overall task performance, with average earnings of £36. We used all 29 participants for behavioral analyses and model fits. We excluded trials without a response within the 3-s deadline (175 trials in total; median 3 per participant, range 0-86) from all analyses. A physician identified one participant’s brain as abnormal. Three participants exhibited excessive head motion during the task (more than 3 mm in any direction). We excluded imaging data from these four participants, leaving 25 for the fMRI analyses. The University of Birmingham Ethics Board approved the experiment (program ID ERN_20-1897AP11), and all subjects provided written informed consent.

### Procedures

Each experimental session lasted less than 2 hours, including 30 minutes of document procedures and training, 10 minutes for scanner setup, 1 hour in the scanner, and 10 minutes of post-scanner procedures. Before entering the scanner, participants completed the consent forms and took a 15-minute tutorial to learn the task and practice 2 games. During behavioral training, subjects used two arrow keys on a laptop keyboard (left and right). Participants were then changed into MR-safe clothes, given earplugs, screened for metal, and positioned in the 3T Siemens MRI scanner. After completing the task, participants saw their total earnings before leaving the lab.

### Task

We programmed the task in MATLAB using Psychtoolbox-3 (http://psychtoolbox.org/). Each session included three scanner runs. Each run consisted of 10 games, and each game included 10 choice trials (resulting in 300 trials per participant). Before the first trial of each run (1, 2, and 3), the computer displayed a “GET READY” screen for 5 seconds and waited for a scanner trigger to begin the run.

In each game, a participant played with a new 20-card deck made up of four standard suits (hearts, clubs, spades, and diamonds) and five digits (2 to 6), with exactly one card for each suit-digit combination. At the bottom of the screen, the computer displayed a full grid showing the entire deck and highlighted the two current options (**Figure 1**). On each trial, the computer randomly drew two cards without replacement from the remaining deck and presented them as large stimuli on the left and right sides of the top part of the screen, while also indicating their positions in the grid.

Participants pressed the keys on the button box with their index and middle fingers to select the corresponding card (left or right) within a 3-second deadline. We defined response time (RT) as the interval from stimulus onset to keypress. After a valid choice, we highlighted the selected option and kept the display for a duration equal to 1 second plus the remaining time in the 3-second response window (i.e., feedback duration = 1 second + [3 seconds – RT]), ensuring that the choice and feedback periods were the same across trials. The chosen card was then highlighted in white, and the unchosen card was removed from both the choice set and the grid.

If participants missed the deadline, the computer removed both cards, displayed “TOO LATE” for 1 second, and assigned zero points for that trial. We jittered the inter-trial interval from 2 to 6 seconds using a pre-generated schedule that was independently shuffled for each of the three runs.

Participants were given full instructions on how their bonuses were calculated. The participants scored each selected card based on its digit value (2 to 6 points). After each game (10 trials), the computer calculated and displayed the round total and awarded two set bonuses. It also tallied suit sets and digit sets of at least four cards, giving 20 points for each suit set and each digit set, then displayed the round summary and cumulative set counts. Throughout the session, the computer tracked the total number of suit sets and digit sets. It then calculated a session bonus by multiplying 20 points by the larger of these two cumulative counts.

### Summary of the task

(1) Each game started with a fresh, fully visible deck of 20 cards that included four suits × five digits (2-6), one card of each combination. (2) On each of the 10 rounds, two cards were drawn at random from the remaining deck. (3) The participant chose one card and earned its digit value in points immediately. The other card was discarded for the rest of the game. (4) After round 10, the deck was exhausted and the game ended. Each suit set (any four cards of the same suit) and each digit set (all four cards of the same digit) earned the participant 20 bonus points. A kept card could count toward one suit set and one digit set at the same time. (5) At the end of the whole session, a further bonus of 20 points was paid for every set of the more frequently completed type (suit or digit) across all 30 games. Throughout the game, the grid showed which cards had been kept, discarded, or were still in the deck, so participants could always see which sets remained possible and rely less on working memory.

We calculated the final score as the sum of round totals plus the total session bonus. We converted points to a monetary bonus at a rate of 500 points per £1 and paid a £30 show-up fee in addition to any performance-based bonus.

### Imaging methods

We acquired functional images using a 3T Siemens Prisma scanner with a 32-channel head coil at the Centre for Human Brain Health (University of Birmingham). We obtained a T1-weighted anatomical scan and three runs, each with 700 acquisitions.

For the structural scan, we used a 3D inversion-recovery prepared gradient-echo sequence (Siemens MPRAGE) with these parameters: TR = 2.0 s, TE = 2.03 ms, TI = 880 ms, flip angle = 8°, GRAPPA = 2 with 24 reference lines, bandwidth = 240 Hz/pixel, phase oversampling = 5%, base matrix = 256 × 256 (phase-encoding steps = 269), slice/partition thickness = 1.0 mm.

For the functional scans, we used the CMRR multiband gradient-echo EPI sequence (multiband factor = 3; GRAPPA = 2) with 60 oblique slices (thickness = 2.4 mm; no gap; obliquity ≈ –40°). Imaging parameters included TR = 1.254 s, TE = 30 ms, flip angle = 65°, bandwidth = 1735 Hz/pixel, effective echo spacing = 0.33 ms, and total readout time = 29.37 ms.

We preprocessed the data with SPM12 (Wellcome Centre for Human Neuroimaging). First, we performed rigid-body realignment (Estimate & Reslice; two-pass register-to-mean) and saved the resliced images along with the mean functional image. Next, we applied slice-timing correction to the resliced images using the study’s slice order, TR, TA, and the specified reference slice. Then, we co-registered the mean EPI (source) to the T1-weighted structural image (reference) with normalized mutual information, and we ran SPM’s Unified Segmentation on the T1 to compute forward and inverse deformation fields. We normalized the slice-timing-corrected functional images to MNI space by applying the forward deformation field, resampling to 3×3×3 mm within the standard MNI bounding box, and finally smoothing the normalized images with a 6 mm FWHM Gaussian kernel.

As the vmPFC is prone to susceptibility-related signal loss, we computed temporal signal-to-noise ratio (tSNR) maps for each participant from the unsmoothed normalized images (voxel mean divided by the standard deviation after linear detrending, averaged over runs). Mean tSNR was 39-71 in the seven regions of interest (vmPFC 49 ± 6, minimum 38 across participants), and every ROI voxel was inside the analysis mask of every participant (**Supplementary Figure S7**; **Supplementary Table S4**).

### Computational modeling

We used 8 models of how participants value the two cards offered on each trial. All models shared the same choice rule, the binary logit (softmax) rule. Specifically, the probability that the participant keeps the right card (*R*) rather than the left card (*L*) is a logistic function of the difference between the two cards’ utilities,

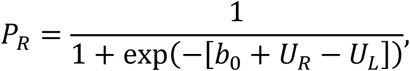

where *P_R_* is the probability of choosing the right card, *U_R_* and *U_L_* are the subjective utilities of the right and left card, computed by each model as described below, and *b*_0_ is a side bias (*b*_0_ > 0 means a tendency to press the right button). Utilities are in arbitrary units whose scale absorbs the softmax inverse temperature; the larger the weights, the more deterministically choices follow utility differences, and the ratio of two weights is independent of this scale.

### Card variables

For a card *c* in a given trial, the models use the following quantities, all computed from the cards the participant has already retained, the cards still in the deck, and the two cards presented as choice options:

*V_c_* is the point value of card *c* (its digit, 2-6), earned immediately if the card is chosen.

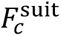 equals 1 if keeping card *c* would leave its suit set still achievable; otherwise it equals 0. The other presented card is not counted for this, because it is discarded if *c* is kept.

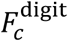 equals 1 if that digit set remains achievable if the card is kept; otherwise it equals 0.

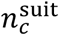 and 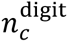 are the numbers of cards of card *c*’s suit and of its digit that the participant has already kept in the current game (0-3); they measure progress toward each set.

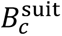 and 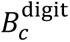 are the expected bonus points (20 points per completed set) from suit sets and from digit sets that would be completed between the current choice and the end of the game if card *c* is kept and the other card discarded, and 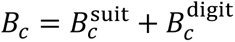 is the total expected bonus. 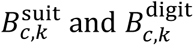 are the parts of these expectations that come from sets completed exactly *k* rounds after the current choice (*k* = 0 implies the set is completed by card *c* itself). W*_c_* is the expected number of points from the cards kept in the remaining rounds. These expectations are estimated by simulating the rest of the game (see the planning models below) and are used only by M4-M8.

### Models

Each model defines the utility *U_c_* of a card from these variables. All weights are free parameters estimated separately for each participant, together with the side bias *b*_0_.

### M1 (myopic)

The baseline model values a card only by its immediate points:

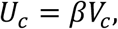

where *β* (≥ 0) is the value weight, the utility of one immediate point. Free parameters: *b*_0_, *β*.

### M2 (feasibility heuristic)

A card gains a fixed amount of value for each set it could still help complete. This is a simple cue-based heuristic that ignores both the probability of completing the set and how far the participant has progressed toward it:

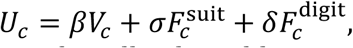

where *σ* and *δ* are the utilities of a still-achievable suit set and digit set. Their ratios to the value weight, σ/β and δ/β, are the numbers of immediate points a participant would give up to keep a suit or digit set achievable. Free parameters: *b*_0_, *β*, *σ*, *δ*.

### M3 (goal-gradient heuristic)

In this model, the value of an achievable set grows with the progress already made toward it. We used a power functional form but also fitted hyperbolic and exponential functions of the number of cards still needed (see the Robustness section below):

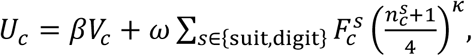

where the sum runs over the card’s two sets (*s* = suit, digit), 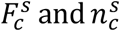 are the feasibility indicator and progress count defined above, and 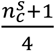 is the fraction of the four-card set that would be in hand if card *c* were kept (from 1/4 to 1). *ω* (≥ 0) is the utility of an achievable set that card *c* would complete, and *κ* (≥ 0) sets how steeply set value rises with progress; with *κ* = 0 every achievable set is worth *ω* (identical to M2 with σ = δ), and with *κ* = 2 a set of which three cards are already held is worth sixteen times more than an untouched set. Free parameters: *b*_0_, *β*, *ω*, *κ* (to restrict the parameter space, к is estimated over 0, 0.5, 1, 1.5, 2, 3, 4).

### M4-M8 (planning models)

These models compute prospective value by forward simulation, using the rollout method. The value of an action is estimated by simulating many continuations of the problem in which later decisions follow a fixed, reasonable base policy^74,75^. For each card offered on a trial, the model simulates 2,000 continuations of the game from the current state: the card is kept, the other card is discarded, and each remaining round draws two random cards from the deck. Choices in the simulated future follow the best-scoring simple policy we identified (immediate points plus 5 for each achievable set; this policy achieves 83.7 points per game in simulation, compared with the myopic policy of always taking the higher card, which achieves 74.3 points). Averaging over the 2,000 simulations produces 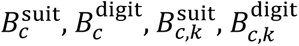 and W*c*.

### M4 (planner)

The card is valued by its immediate points and the bonus it is expected to lead to:

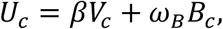

where *ω_B_* (≥ 0) is the utility of one expected bonus point. The ratio *ω_B_*/*β* is the participant’s exchange rate between expected future bonus points and immediate points; an agent that maximizes expected points has *ω_B_*/*β* = 1. Free parameters: *b*_0_, *β*, *ω_B_*.

### M5 (planner with separate set types)

Suit and digit bonuses have separate weights, which allows for the end-of-session bonus making one set type worth more than the other:

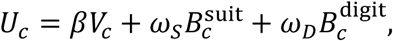

where *ω_S_* and *ω_D_* are the utilities of one expected bonus point from suit sets and from digit sets. *ω_S_* > *ω_D_* would indicate that suit sets are valued more. Free parameters: *b*_0_, *β*, *ω_S_*, *ω_D_*.

### M6 (discounting planner)

This model assumes participants plan only a limited distance ahead and give less weight to outcomes deep in the decision tree. It therefore discounts bonuses from sets completed further in the future:

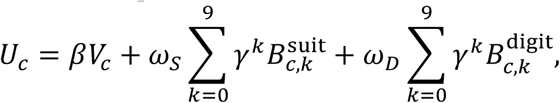

where *γ* (0 ≤ y ≤ 1) is a per-round discount factor, so a bonus from a set completed *k* rounds after the current choice is weighted by *γ^k^*; *ω_S_* and *ω_D_* are as in M5. With *γ* = 1 the model reduces to M5, and with *γ* = 0 only sets completed by card *c* itself count. Free parameters: *b*_0_, *β*, *ω_S_*, *ω_D_*, *γ* (y estimated over 0, 0.1, …, 1 for tractability).

### M7 (normative Q-value)

The card is valued by its action value, the total number of points it is expected to yield over the rest of the game^76^ so that immediate and future points are traded one for one:

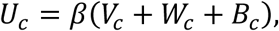

where *V_c_* + W*_c_* + *B_c_* is the expected total of immediate points, future card points and set bonuses, and *β* is the only weight (the inverse temperature). Free parameters: *b*_0_, *β*.

### M8 (hybrid)

Choices could combine a simple heuristic with planning, as in hybrid accounts of model-free and model-based control^46^. In short, M8 combines the feasibility heuristic M2 with the planner M5 with separate set types:

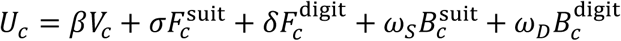

with all parameters as defined for M2 and M5. Free parameters: *b*_0_, *β*, *σ*, *δ*, *ω_S_*, *ω_D_*.

### Robustness to the form of the goal gradient

For M3, we fitted two alternatives in which the set value falls with the number of cards *r* still needed to complete the set: a hyperbolic gradient, 1/(1 + ℎ · *r*), and an exponential gradient, *exp*(−*λ* · *r*), each with one steepness parameter as in M3. The power form (M3) fit best of the three (summed BIC 43 lower than the exponential and 124 lower than the hyperbolic form; protected exceedance probability > 0.99), and all three reproduced the progress effect on conflict trials (predicted probability of choosing the set card with three cards already held: 0.71-0.72; observed 0.74). The three forms were only partly distinguishable in model recovery (correctly recovered in 79%, 48% and 22% of simulations; **Supplementary Figure S9**).

### Model fitting and comparison

We fitted every model to each participant using maximum likelihood estimation (MLE). We compared models by the summed Bayesian information criterion (BIC), by random-effects Bayesian model selection on –BIC/2^77^ reporting protected exceedance probabilities^78^, and by leave-one-run-out cross-validated log likelihood and balanced accuracy. Mixed-effects logistic regression versions of M1 and M2 are reported in **Supplementary Table S5**.

### Model and parameter recovery

For each model, we simulated 116 synthetic participants (four per real participant, using that participant’s fitted parameters). We recomputed all model variables on the simulated trials, fitted all eight models to every synthetic data set, and computed how often each generating model was best fit by BIC.

### Behavioral analyses

We classified trials as congruent (the higher-point card also had the higher feasible set value), conflict (the higher-point card had the lower set value), or neutral (equal points or equal set value). We analyzed response times (RT) using linear mixed models of log RT that included trial-within-game and session trial as covariates and participant random slopes. To avoid circularity in parameter-performance relationships, we fitted the models to the odd-numbered games only and correlated the parameters with points earned in the even-numbered games, and vice versa. All analyses used R 4.4 with lme4 1.1-35.5.

### Model-based regressors for fMRI

We produced trial-wise regressors from the winning goal-gradient model (M3). We fixed the value and set weights at their group medians (β = 0.66, ω = 2.31), so that every participant’s regressors were on the same utility scale and individual differences in choice consistency did not enter the neural estimates. In GLMs 1, 2 and 4, prospective utility followed each participant’s own gradient steepness к (range 0-4, median 2 in the fMRI sample). For each trial, we computed the immediate utility of the chosen card (βV, proportional to its points), its prospective utility (the set term of M3), the total utility of the unchosen card, the signed decision value (utility of the chosen minus the unchosen card), and its absolute value. For the planner, we used the chosen card’s points and its expected set bonus B (M4).

### fMRI analysis

#### First-level models

We estimated first-level general linear models (GLMs) in SPM25 (Wellcome Centre for Human Neuroimaging). We included the three runs in one model as separate sessions, with a 128-s high-pass filter and the FAST model of serial correlations. We modeled each choice as an event at stimulus onset with a duration equal to the response time, convolved with the canonical hemodynamic response function. We mean-centered parametric modulators within each run and entered them without orthogonalization, so that each effect was estimated while controlling for all others. Each model-based GLM also included four controls (trial within the game, trial within the session, response time, and response button) and the six realignment parameters of each run. We did not include no-response trials. We excluded one participant’s run as it was incomplete.

#### GLMs

We tested five GLMs (**Supplementary Table S6**).

GLM1 (value components) contained the immediate utility of the chosen card, its prospective utility, and the utility of the unchosen card. We contrasted each regressor against 0, prospective minus immediate utility, and chosen minus unchosen utility. Correlations between the three regressors were low (|r| ≤ 0.19; all variance inflation factors < 1.4). The unchosen card value was a single total utility.

GLM2 (decision value) included the signed decision value and its absolute value.

GLM3 (conflict) modeled trials as separate conditions by trial type (congruent, neutral, conflict) and by whether the lower-point card was chosen, with ties as a further condition and no parametric modulators. This GLM tested whether choosing the lower-point card engages different regions when prospective value supports it (conflict trials) than when it does not (neutral trials).

GLMs 4 and 5 included the immediate and prospective components of the chosen card’s value, computed either by the goal-gradient model (GLM 4: immediate and prospective utility) or by the planner (GLM 5: points and expected set bonus). They differed only in how prospective value was computed and were compared with cross-validated Bayesian model selection (see below).

#### Group inference

We used sign-flipping permutation tests (5,000 permutations) on variance-smoothed pseudo-t statistics (6-mm FWHM), the one-sample procedure of SnPM^79^, with a cluster-forming threshold of p < 0.001 (|t| ≥ 3.47, 24 df) and cluster-level family-wise error (FWE) control at p < 0.05. We report all clusters that survived FWE correction (**Supplementary Table S2**). We labeled the peaks with the Neuromorphometrics atlas distributed with SPM. We displayed the maps with MRIcroGL (https://www.nitrc.org/projects/mricrogl).

#### Regions of interest

We used seven regions of interest (ROIs) defined independently of our data (**Supplementary Figure S10**): the vmPFC and striatal clusters from a meta-analysis of subjective-value signals^15^, and the left and right caudate, left and right insula and precuneus from the AAL atlas. For each participant and contrast, we averaged the contrast estimates over the voxels of each ROI and tested them against zero with two-sided one-sample t-tests, Bonferroni-corrected over the seven ROIs. To quantify evidence for the absence of an effect, we also computed default JZS Bayes factors (Cauchy prior scale √2/2^80^); BF4o < 1/3 is moderate evidence for no effect and BF4o > 3 is moderate evidence for an effect. We included the precuneus, as the posterior medial cortex is consistently activated by subjective value^15,16^.

#### Neural model comparison

To ask whether neural prospective value is better described by the goal-gradient heuristic or by the planner, we compared GLMs 4 and 5 using cross-validated Bayesian model selection in the MACS toolbox^81,82^. For each participant and voxel, MACS computes the cross-validated log model evidence of each GLM, using each run in turn as test data. We averaged these values within each ROI and summarized the comparison by the share of participants with stronger evidence for GLM 4, the expected model frequency from random-effects Bayesian model selection^77^ and the summed log Bayes factor (**Supplementary Table S3**).

## Data availability

The behavioral data and unthresholded and thresholded group-level statistical maps for all contrasts are available on Zenodo (https://doi.org/10.5281/zenodo.23980339). The individual fMRI data are available from the corresponding author upon request.

## Code availability

The analysis code is available on Zenodo (https://doi.org/10.5281/zenodo.23980339).

## Generative AI

We used Claude Opus 5.5 for coding assistance and partial summarization of methods and results, and verified all outputs.

## Acknowledgments

We thank Romy Froemer for helpful feedback and conversations.

## Author contributions

A.K. designed the experiment and analyses. A.K. programmed the experiment. D.L. and A.K. collected the data. A.K. performed the data analysis and wrote the initial draft of the paper. D.L. co-wrote the paper. A.K. supervised the project.

## REFERENCES

1. Mattar, M. G. & Lengyel, M. Planning in the brain. Neuron 110, 914–934 (2022).

2. Balaguer, J., Spiers, H., Hassabis, D. & Summerfield, C. Neural Mechanisms of Hierarchical Planning in a Virtual Subway Network. Neuron 90, 893–903 (2016).

3. Kuperwajs, I., Russek, E. M., Mattar, M. G., Ma, W. J. & Griffiths, T. L. Looking deeper into the algorithms underlying human planning. Trends Cogn. Sci. 0, (2025).

4. Simon, D. A. & Daw, N. D. Neural Correlates of Forward Planning in a Spatial Decision Task in Humans. J. Neurosci. 31, 5526–5539 (2011).

5. Jensen, K. T., Hennequin, G. & Mattar, M. G. A recurrent network model of planning explains hippocampal replay and human behavior. Nat. Neurosci. 27, 1340–1348 (2024).

6. Kuperwajs, I., Schütt, H. H. & Ma, W. J. Using deep neural networks as a guide for modeling human planning. Sci. Rep. 13, 20269 (2023).

7. van Opheusden, B. et al. Expertise increases planning depth in human gameplay. Nature 618, 1000–1005 (2023).

8. Ingvar, D. H. ‘ Memory of the future’: an essay on the temporal organization of conscious awareness. Hum. Neurobiol. 4, 127–136 (1985).

9. Suddendorf, T. & Corballis, M. C. The evolution of foresight: What is mental time travel, and is it unique to humans? Behav. Brain Sci. 30, 299–313 (2007).

10. Shallice, T. Specific impairments of planning. Philos. Trans. R. Soc. Lond. B Biol. Sci. 298, 199–209 (1982).

11. Passini, R., Rainville, C., Marchand, N. & Joanette, Y. Wayfinding in dementia of the Alzheimer type: Planning abilities. J. Clin. Exp. Neuropsychol. 17, 820–832 (1995).

12. Valls-Serrano, C., Verdejo-García, A. & Caracuel, A. Planning deficits in polysubstance dependent users: Differential associations with severity of drug use and intelligence. Drug Alcohol Depend. 162, 72–78 (2016).

13. Kable, J. W. & Glimcher, P. W. The neural correlates of subjective value during intertemporal choice. Nat. Neurosci. 10, 1625–1633 (2007).

14. McClure, S. M. Separate Neural Systems Value Immediate and Delayed Monetary Rewards. Science 306, 503–507 (2004).

15. Bartra, O., McGuire, J. T. & Kable, J. W. The valuation system: A coordinate-based meta-analysis of BOLD fMRI experiments examining neural correlates of subjective value. NeuroImage 76, 412–427 (2013).

16. Clithero, J. A. & Rangel, A. Informatic parcellation of the network involved in the computation of subjective value. Soc. Cogn. Affect. Neurosci. 9, 1289–1302 (2014).

17. Levy, D. J. & Glimcher, P. Comparing apples and oranges: Using reward-specific and reward-general subjective value representation in the brain. J. Neurosci. 31, 14693–14707 (2011).

18. Peters, J. & Büchel, C. Neural representations of subjective reward value. Behav. Brain Res. 213, 135–141 (2010).

19. Cox, K. M. & Kable, J. W. BOLD subjective value signals exhibit robust range adaptation. J. Neurosci. 34, 16533–16543 (2014).

20. O’Doherty, J. P.. Cockburn, J. & Pauli,W. M. Learning, Reward, and Decision Making. Annu. Rev. Psychol. 68, 73–100 (2017).

21. McClure, S. M., Ericson, K. M., Laibson, D. I., Loewenstein, G. & Cohen, J. D. Time Discounting for Primary Rewards. J. Neurosci. 27, 5796–5804 (2007).

22. Holton, E. et al. Disentangling the Component Processes in Complex Planning Impairments Following Ventromedial Prefrontal Lesions. J. Neurosci. 45, (2025).

23. Peters, J. & Büchel, C. Episodic future thinking reduces reward delay discounting through an enhancement of prefrontal-mediotemporal interactions. Neuron 66, 138–148 (2010).

24. Shenhav, A., Botvinick, M. M. & Cohen, J. D. The expected value of control: an integrative theory of anterior cingulate cortex function. Neuron 79, 217–240 (2013).

25. Kennerley, S. W., Walton, M. E., Behrens, T. E. J., Buckley, M. J. & Rushworth, M. F. S. Optimal decision making and the anterior cingulate cortex. Nat. Neurosci. 9, 940–947 (2006).

26. Cowen, S. L., Davis, G. A. & Nitz, D. A. Anterior cingulate neurons in the rat map anticipated effort and reward to their associated action sequences. J. Neurophysiol. 107, 2393–2407 (2012).

27. Kolling, N., Behrens, T. E. J., Mars, R. B. & Rushworth, M. F. S. Neural mechanisms of foraging. Science 336, 95–98 (2012).

28. Figner, B. et al. Lateral prefrontal cortex and self-control in intertemporal choice. Nat. Neurosci. 13, 538–539 (2010).

29. Knutson, B., Adams, C. M., Fong, G. W. & Hommer, D. Anticipation of increasing monetary reward selectively recruits nucleus accumbens. J. Neurosci. 21, RC159 (2001).

30. Haber, S. N. & Knutson, B. The reward circuit: Linking primate anatomy and human imaging. Neuropsychopharmacology 35, 4–26 (2010).

31. O’Doherty, J. et al. Dissociable roles of ventral and dorsal striatum in instrumental conditioning. Science 304, 452–454 (2004).

32. Balleine, B. W., Delgado, M. R. & Hikosaka, O. The role of the dorsal striatum in reward and decision-making. J. Neurosci. 27, 8161–8165 (2007).

33. Wunderlich, K., Smittenaar, P. & Dolan, R. J. Dopamine Enhances Model-Based over Model-Free Choice Behavior. Neuron 75, 418–424 (2012).

34. Howe, M. W., Tierney, P. L., Sandberg, S. G., Phillips, P. E. & Graybiel, A. M. Prolonged dopamine signalling in striatum signals proximity and value of distant rewards. Nature 500, 575–579 (2013).

35. Hamid, A. A. et al. Mesolimbic dopamine signals the value of work. Nat. Neurosci. 19, 117–126 (2015).

36. Mohebi, A. et al. Dissociable dopamine dynamics for learning and motivation. Nature 570, 65–70 (2019).

37. Kuhnen, C. M. & Knutson, B. The neural basis of financial risk taking. Neuron 47, 763–770 (2005).

38. Preuschoff, K., Quartz, S. R. & Bossaerts, P. Human Insula Activation Reflects Risk Prediction Errors As Well As Risk. J. Neurosci. 28, 2745–2752 (2008).

39. Knutson, B., Rick, S., Wimmer, G. E., Prelec, D. & Loewenstein, G. Neural predictors of purchases. Neuron 53, 147–156 (2007).

40. Sanfey, A. G., Rilling, J. K., Aronson, J. A., Nystrom, L. E. & Cohen, J. D. The neural basis of economic decision-making in the Ultimatum Game. Science 300, 1755–1758 (2003).

41. Craig, A. D. How do you feel — now? The anterior insula and human awareness. Nat. Rev. Neurosci. 10, 59–70 (2009).

42. Menon, V. & Uddin, L. Q. Saliency, switching, attention and control: a network model of insula function. Brain Struct. Funct. 214, 655–667 (2010).

43. Keramati, M., Dezfouli, A. & Piray, P. Speed/Accuracy Trade-Off between the Habitual and the Goal-Directed Processes. PLoS Comput. Biol. 7, e1002055 (2011).

44. Keramati, M., Smittenaar, P., Dolan, R. J. & Dayan, P. Adaptive integration of habits into depth-limited planning defines a habitual-goal-directed spectrum. Proc. Natl. Acad. Sci. 113, 12868–12873 (2016).

45. Sezener, C. E., Dezfouli, A. & Keramati, M. Optimizing the depth and the direction of prospective planning using information values. PLOS Comput. Biol. 15, e1006827 (2019).

46. Daw, N. D., Gershman, S. J., Seymour, B., Dayan, P. & Dolan, R. J. Model-based influences on humans’ choices and striatal prediction errors. Neuron 69, 1204–1215 (2011).

47. Daw, N. D. Model-based reinforcement learning as cognitive search: neurocomputational theories. Cogn. Search Evol. Algorithms Brain http://citeseerx.ist.psu.edu/viewdoc/download?rep=rep1&type=pdf&doi=10.1.1.216.209 (2012).

48. Daw, N. D. Are we of two minds? Nat. Neurosci. 21, 1497 (2018).

49. Daw, N. D. & Dayan, P. The algorithmic anatomy of model-based evaluation. Philos. Trans. R. Soc. B Biol. Sci. 369, 20130478–20130478 (2014).

50. Bornstein, A. M. & Daw, N. D. Cortical and Hippocampal Correlates of Deliberation during Model-Based Decisions for Rewards in Humans. PLoS Comput. Biol. 9, e1003387 (2013).

51. Bornstein, A. M. & Daw, N. D. Dissociating hippocampal and striatal contributions to sequential prediction learning: Sequential predictions in hippocampus and striatum. Eur. J. Neurosci. 35, 1011–1023 (2012).

52. Konovalov, A. & Krajbich, I. Neurocomputational Dynamics of Sequence Learning. Neuron 98, 1282–1293.e4 (2018).

53. Russek, E. M., Momennejad, I., Botvinick, M. M., Gershman, S. J. & Daw, N. D. Predictive representations can link model-based reinforcement learning to model-free mechanisms. PLOS Comput. Biol. 13, e1005768 (2017).

54. Momennejad, I., Otto, A. R., Daw, N. D. & Norman, K. A. Offline replay supports planning in human reinforcement learning. eLife 7, e32548 (2018).

55. Konovalov, A. & Krajbich, I. Gaze data reveal distinct choice processes underlying model-based and model-free reinforcement learning. Nat. Commun. 7, 12438 (2016).

56. Konovalov, A. & Krajbich, I. Mouse tracking reveals structure knowledge in the absence of model-based choice. Nat. Commun. 11, 1893 (2020).

57. Doll, B. B., Duncan, K. D., Simon, D. A., Shohamy, D. & Daw, N. D. Model-based choices involve prospective neural activity. Nat. Neurosci. https://doi.org/10.1038/nn.3981 (2015) doi:10.1038/nn.3981.

58. Hare, T. A., O’Doherty, J., Camerer, C. F., Schultz, W. & Rangel, A. Dissociating the Role of the Orbitofrontal Cortex and the Striatum in the Computation of Goal Values and Prediction Errors. J. Neurosci. 28, 5623–5630 (2008).

59. Feher da Silva, C., Lombardi, G., Edelson, M. & Hare, T. A. Rethinking model-based and model-free influences on mental effort and striatal prediction errors. Nat. Hum. Behav. 1–14 (2023) doi:10.1038/s41562-023-01573-1.

60. Moneta, N., Garvert, M. M., Heekeren, H. R. & Schuck, N. W. Task state representations in vmPFC mediate relevant and irrelevant value signals and their behavioral influence. Nat. Commun. 14, 3156 (2023).

61. Garvert, M. M., Saanum, T., Schulz, E., Schuck, N. W. & Doeller, C. F. Hippocampal spatio-predictive cognitive maps adaptively guide reward generalization. Nat. Neurosci. 1–12 (2023) doi:10.1038/s41593-023-01283-x.

62. Schuck, N. W., Cai, M. B., Wilson, R. C. & Niv, Y. Human Orbitofrontal Cortex Represents a Cognitive Map of State Space. Neuron 91, 1402–1412 (2016).

63. Gershman, S. J. & Daw, N. D. Reinforcement learning and episodic memory in humans and animals: an integrative framework. Annu. Rev. Psychol. 68, 101–128 (2017).

64. Lee, S. W., Shimojo, S. & O’Doherty, J. P. Neural Computations Underlying Arbitration between Model-Based and Model-free Learning. Neuron 81, 687–699 (2014).

65. Hull, C. L. The goal-gradient hypothesis and maze learning. Psychol. Rev. 39, 25–43 (1932).

66. Kivetz, R., Urminsky, O. & Zheng, Y. The goal-gradient hypothesis resurrected: Purchase acceleration, illusionary goal progress, and customer retention. J. Mark. Res. 43, 39–58 (2006).

67. Huys, Q. J. M. et al. Bonsai Trees in Your Head: How the Pavlovian System Sculpts Goal-Directed Choices by Pruning Decision Trees. PLoS Comput. Biol. 8, e1002410 (2012).

68. Callaway, F. et al. Rational use of cognitive resources in human planning. *Nat*. Hum. Behav. 6, 1112–1125 (2022).

69. Cogliati Dezza, I., Maher, C. & Sharot, T. People adaptively use information to improve their internal states and external outcomes. Cognition 228, (2022).

70. Paulus, M. P. & Stein, M. B. An insular view of anxiety. Biol. Psychiatry 60, 383–387 (2006).

71. Redish, A. D. The Mind within the Brain: How We Make Decisions and How Those Decisions Go Wrong. (Oxford University Press, 2013).

72. Schultz, W. A Neural Substrate of Prediction and Reward. Science 275, 1593–1599 (1997).

73. Doll, B. B., Simon, D. A. & Daw, N. D. The ubiquity of model-based reinforcement learning. Curr. Opin. Neurobiol. 22, 1075–1081 (2012).

74. 74. Tesauro, G. & Galperin, G. R. On-line policy improvement using Monte-Carlo search. in 1068-1074 (MIT Press, 1997).

75. Bertsekas, D. P., Tsitsiklis, J. N. & Wu, C. Rollout algorithms for combinatorial optimization. J. Heuristics 3, 245–262 (1997).

76. Sutton, R. S. & Barto, A. G. Reinforcement Learning: An Introduction. (MIT Press, Cambridge, MA, 2018).

77. Stephan, K. E., Penny, W. D., Daunizeau, J., Moran, R. J. & Friston, K. J. Bayesian model selection for group studies. NeuroImage 46, 1004–1017 (2009).

78. Rigoux, L., Stephan, K. E., Friston, K. J. & Daunizeau, J. Bayesian model selection for group studies — revisited. NeuroImage 84, 971–985 (2014).

79. Nichols, T. E. & Holmes, A. P. Nonparametric permutation tests for functional neuroimaging: a primer with examples. Hum. Brain Mapp. 15, 1–25 (2002).

80. Rouder, J. N., Speckman, P. L., Sun, D., Morey, R. D. & Iverson, G. Bayesian t tests for accepting and rejecting the null hypothesis. Psychon. Bull. Rev. 16, 225–237 (2009).

81. Soch, J., Haynes, J.-D. & Allefeld, C. How to avoid mismodelling in GLM-based fMRI data analysis: Cross-validated Bayesian model selection. NeuroImage 141, 469–489 (2016).

82. Soch, J. & Allefeld, C. MACS - a new SPM toolbox for model assessment, comparison and selection. J. Neurosci. Methods 306, 19–31 (2018).

