## Supplementary materials for "Striatal Value Signals Track Progress Toward a Goal During Planning"

### SUPPLEMENTARY FIGURES

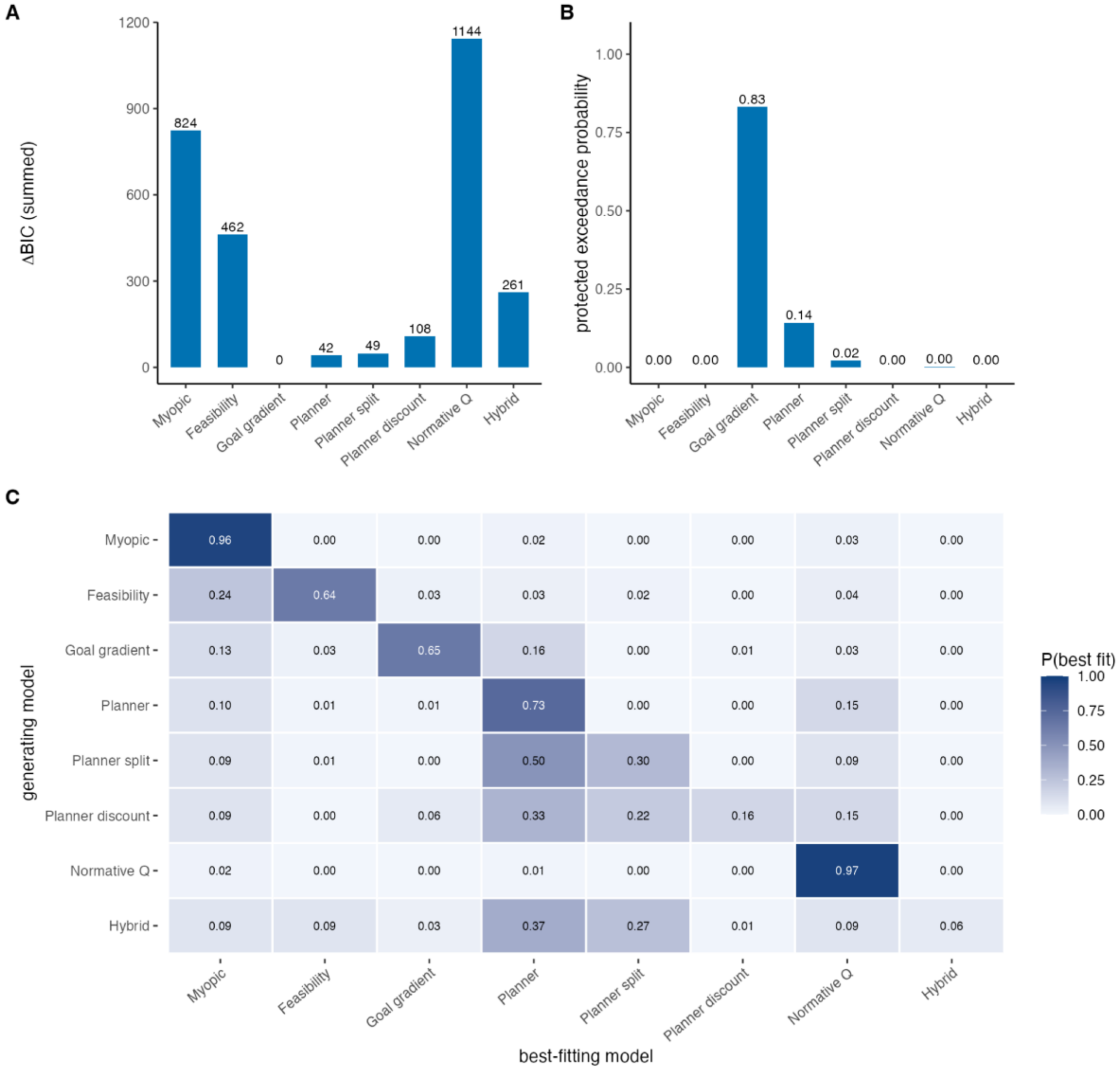


**Supplementary Figure S1. Model comparison and model recovery.** (A) Summed BIC of each model relative to the best model (goal-gradient heuristic, M3). (B) Protected exceedance probabilities from random-effects Bayesian model selection. (C) Model recovery: the share of synthetic data sets generated by each model (rows) that were best fit by each model (columns), 116 synthetic participants per model. Model recovery showed that these conclusions are not artifacts of the comparison procedure. Data generated by the myopic, feasibility, goal-gradient, planner, and normative models were most often best fit by the generating model (96%, 64%, 65%, 73%, and 97%). Data generated by the planner were almost never attributed to the goal-gradient heuristic (1%). N = 29.

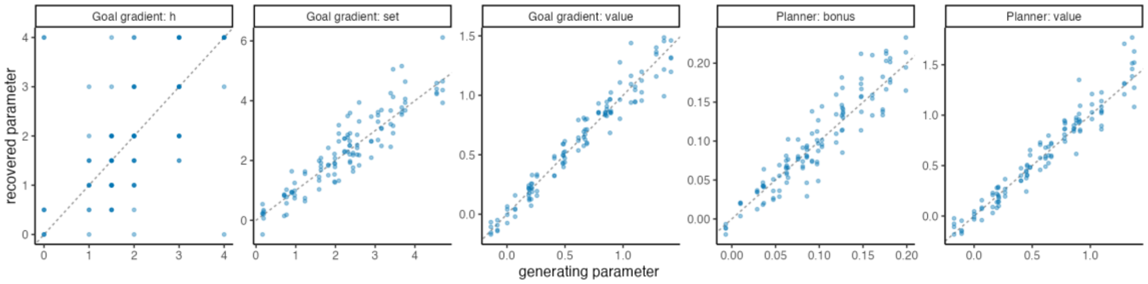


**Supplementary Figure S2. Parameter recovery.** Generating versus recovered parameters for the goal-gradient model (M3: exponent κ, labeled h; set weight ω; value weight β) and the planner (M4: bonus weight ω_B; value weight β), in synthetic data simulated with each participant's fitted parameters (116 synthetic participants per model). The dashed line is the identity line.

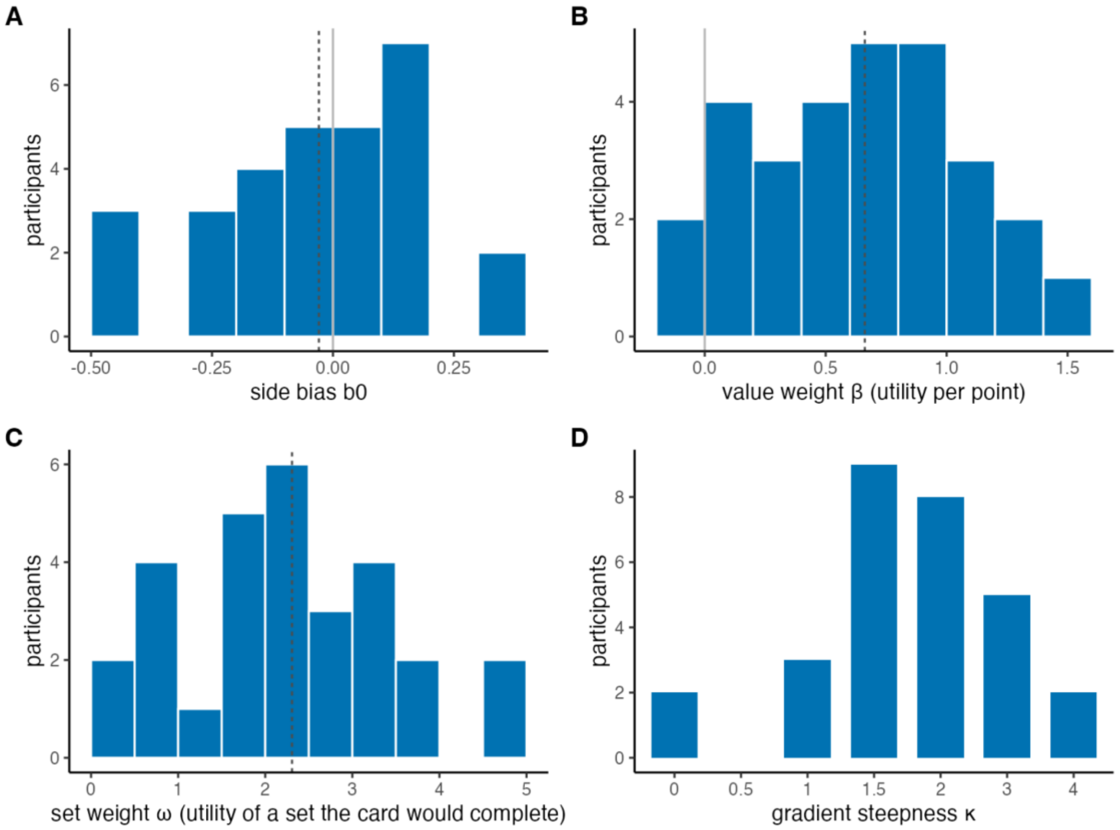


**Supplementary Figure S3. Parameter estimates of the goal-gradient model (M3).** Distributions across participants of (A) the side bias b0, (B) the value weight β, (C) the set weight ω and (D) the gradient steepness κ (estimated over 0, 0.5, 1, 1.5, 2, 3 and 4). Dashed lines mark the medians (b0 = −0.03, β = 0.66, ω = 2.31; the median κ is 2); gray lines mark zero. N = 29.

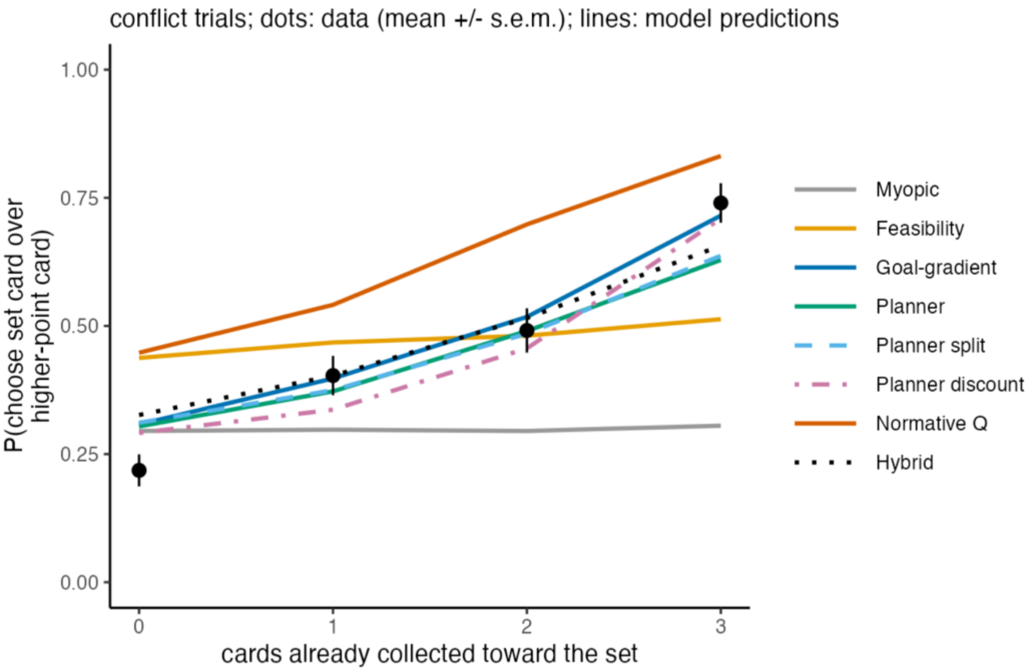


**Supplementary Figure S4. Progress signature on conflict trials for all eight models.** Probability of choosing the set card over the higher-point card as a function of the number of cards of that set already collected; black dots are data (mean ± s.e.m.); lines are the predictions of each model with each participant's fitted parameters. N = 29.

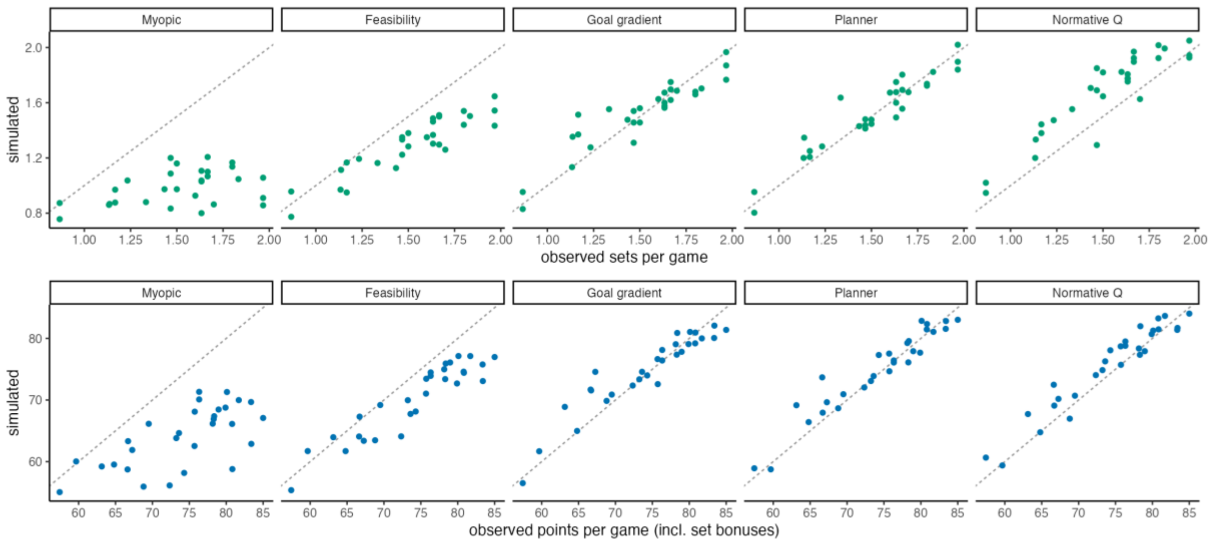


**Supplementary Figure S5. Generative posterior predictive checks.** Observed versus simulated sets per game (top) and points per game including set bonuses (bottom) when each participant's full session was simulated with that participant's fitted parameters, for the myopic, feasibility, goal-gradient, planner, and normative models. Dots are individual participants; the dashed line is the identity line. N = 29.

| | | vmPFC | striatum | L caudate | R caudate | L insula | R insula | precuneus |
|---|---|---|---|---|---|---|---|---|
| Value components | prospective value of the chosen card | 2.4 | **34*** | 6 | **9.6*** | 5 | 0.94 | **62*** |
| | immediate value of the chosen card | 0.24 | 0.24 | 0.24 | 0.21 | **8.2*** | 2.7 | 1.2 |
| | utility of the unchosen card | 0.3 | 0.27 | 0.34 | 0.24 | 0.22 | 0.22 | 0.22 |
| | chosen minus unchosen utility | 0.88 | 1.8 | 0.41 | 1.1 | 0.21 | 0.23 | **24*** |
| | prospective minus immediate value | 1.2 | **14*** | 4.1 | 3.9 | **44*** | 4.4 | 1.5 |
| Decision value | signed decision value (chosen - unchosen) | 0.21 | 0.22 | 0.21 | 0.21 | 1.4 | 0.44 | 0.49 |
| | unsigned decision value | 0.44 | 0.63 | 0.28 | 0.38 | 3.8 | 0.92 | 3.2 |
| Conflict | conflict vs congruent trials | 0.21 | 0.24 | 0.26 | 0.22 | 0.34 | 0.26 | 0.21 |
| | conflict trials: lower vs higher card chosen | 0.36 | 0.4 | 0.24 | 0.21 | 0.42 | 0.24 | 0.22 |
| | lower card chosen: conflict vs neutral | 0.21 | 0.4 | 0.25 | 0.24 | 0.37 | 0.27 | 0.26 |
| | lower choice x conflict | 0.22 | 0.25 | 0.21 | 0.21 | 0.29 | 0.22 | 0.21 |
| Goal-gradient model | prospective value (goal gradient) | 2.9 | **71*** | **24*** | **23*** | **14*** | 1 | **86*** |
| | immediate value (goal gradient) | 0.27 | 0.22 | 0.21 | 0.22 | 6.3 | 4.1 | 0.71 |
| Planner model | expected set bonus (planner) | 0.29 | **31*** | **25*** | **10*** | 0.34 | 0.26 | **>100*** |
| | points (planner) | 0.25 | 0.21 | 0.22 | 0.24 | 5.8 | 4.6 | 0.67 |

t: 4, 2, 0, -2, -4

Cell colour: group t; number: Bayes factor BF10; * and bold: Bonferroni p < 0.05 over 7 ROIs

**Supplementary Figure S6. ROI results for every contrast of GLMs 1–5.** Cell color shows the group t statistic; number is the Bayes factor $BF_{10}$; bold text with a star shows Bonferroni-corrected $p < 0.05$ over seven ROIs. The difficulty contrast of GLM 2 (the negative of the unsigned decision value) is omitted as it mirrors the unsigned decision value. N = 25.

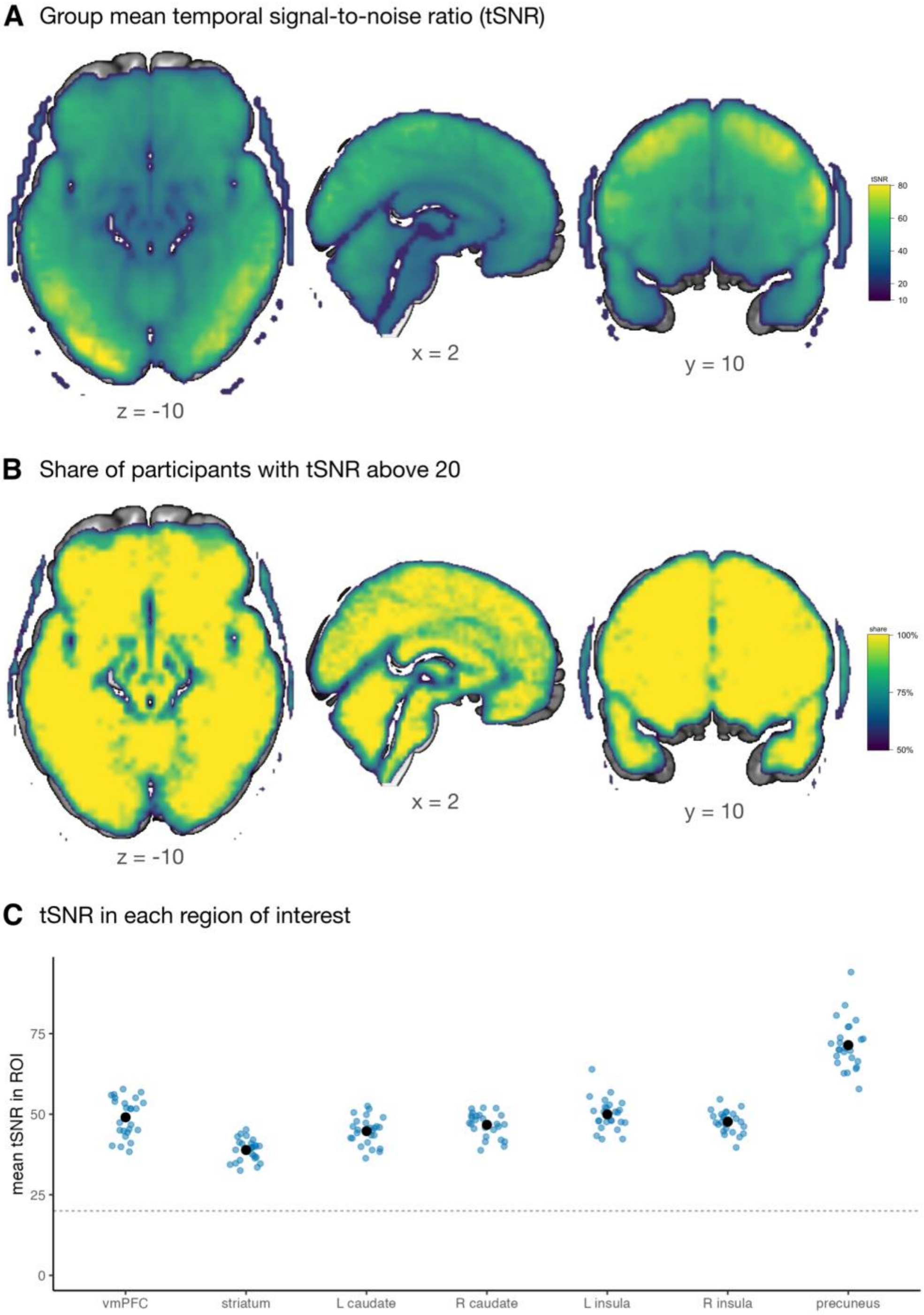


**Supplementary Figure S7. Signal quality.** (A) Group mean temporal signal-to-noise ratio (tSNR), shown in voxels where at least half of the participants had tSNR above 20. (B) Share of participants with tSNR above 20. (C) Mean tSNR in each ROI; small dots are participants; large dots and bars are mean ± s.e.m.; dashed line is tSNR = 20. N = 25.

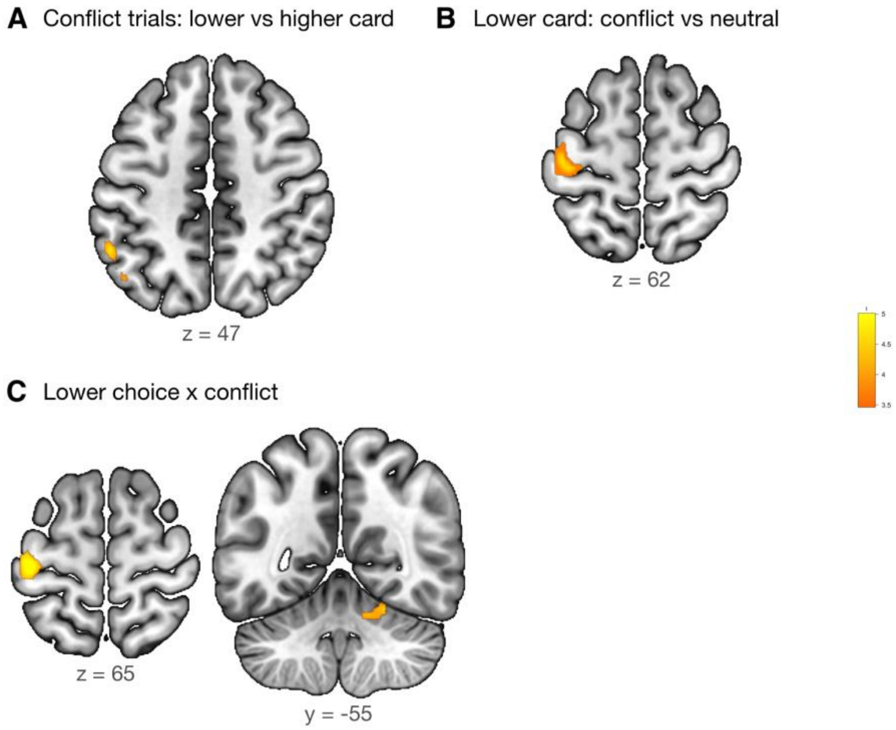


**Supplementary Figure S8. Conflict contrasts (GLM3).** Clusters surviving cluster-level FWE correction ($p < 0.05$; cluster-forming $p < 0.001$) for (A) lower- versus higher-point choices on conflict trials, (B) lower-point choices on conflict versus neutral trials, and (C) the interaction of choice and trial type. The contrast of conflict versus congruent trials produced no surviving clusters. All responses were made with the right hand, so the left sensorimotor and right cerebellar clusters may reflect differences in motor execution between conditions. N = 25.

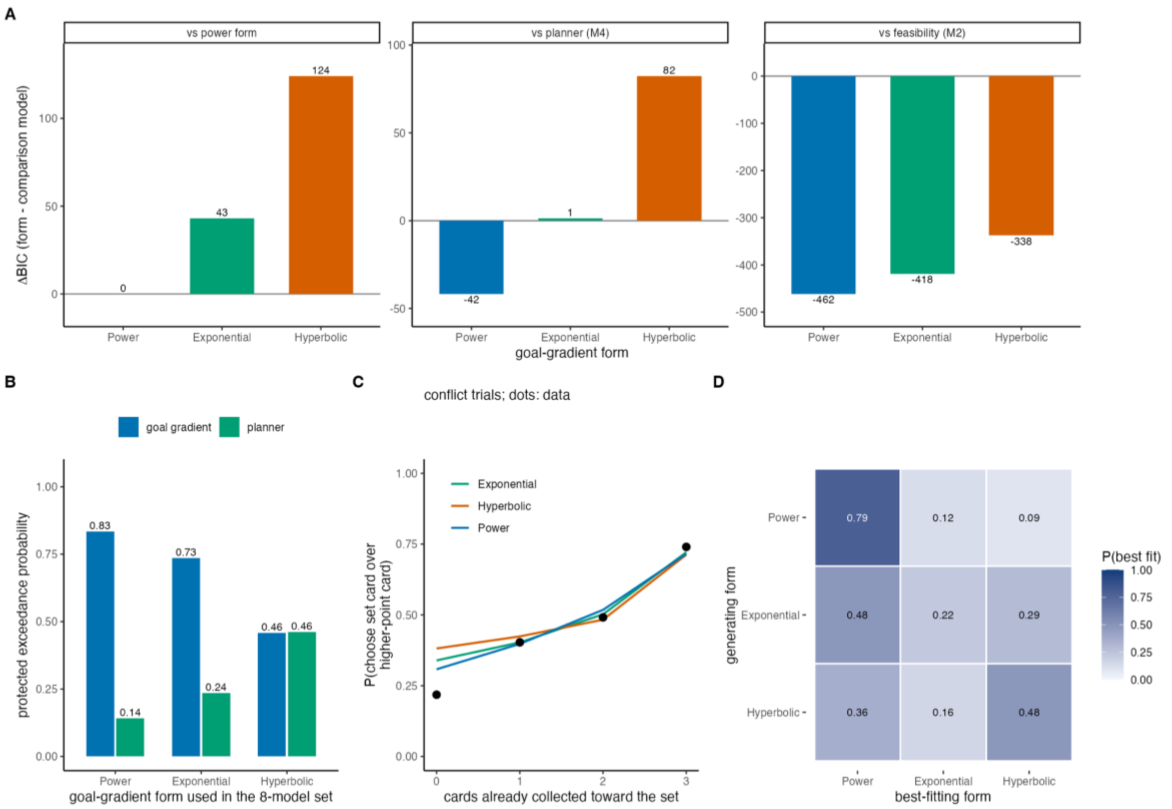


**Supplementary Figure S9. Robustness of the form of the goal gradient.** (A) Summed BIC of the power, exponential, and hyperbolic goal-gradient forms relative to the power form, the planner (M4), and the feasibility heuristic (M2); positive values favor the comparison model. (B) Protected exceedance probabilities of the goal-gradient model and the planner when each form is used in the eight-model comparison. (C) Probability of choosing the set card on conflict trials as a function of progress; dots are the data; lines are the predictions of each form. (D) Model recovery among the three forms. N = 29.

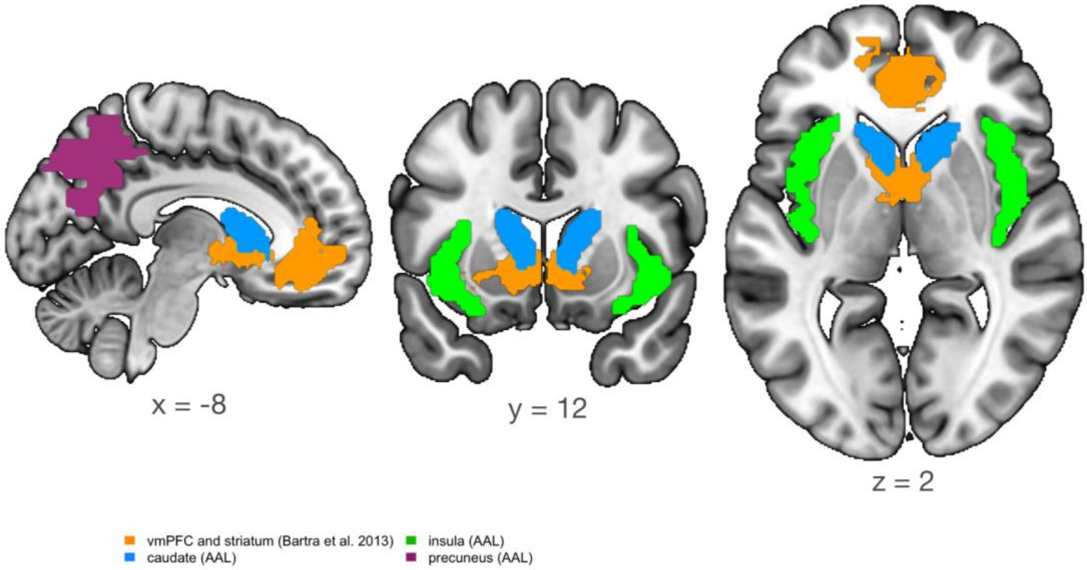


**Supplementary Figure S10. Regions of interest.** The vmPFC and striatal clusters (orange) are from a meta-analysis of subjective-value signals (Bartra et al. 2013); the left and right caudate (blue), left and right insula (green), and precuneus (purple) are from the AAL atlas. All ROIs were defined independently of the data.

### SUPPLEMENTARY TABLES

**Supplementary Table S1. Comparison of the eight computational models.** k is the number of free parameters per participant; ΔBIC is the summed BIC relative to the best model; best fit is the number of participants best fit by BIC; CV log L is the summed leave-one-run-out cross-validated log likelihood; BMS is the expected frequency, and PXP is the protected exceedance probability from random-effects Bayesian model selection. N = 29.

| **Model** | **k** | **Summed BIC** | **ΔBIC** | **Best fit** | **CV log L** | **BMS freq.** | **PXP** |
|---|---|---|---|---|---|---|---|
| M1 Myopic | 2 | 9672 | 824 | 2 | -4836 | 0.06 | 0.001 |
| M2 Feasibility heuristic | 4 | 9310 | 462 | 0 | -4608 | 0.03 | 0.000 |
| M3 Goal-gradient heuristic | 4 | 8848 | 0 | 10 | -4373 | 0.37 | 0.832 |
| M4 Planner | 3 | 8890 | 42 | 7 | -4434 | 0.24 | 0.143 |
| M5 Planner, separate set types | 4 | 8897 | 49 | 5 | -4419 | 0.15 | 0.021 |
| M6 Discounted planner | 5 | 8956 | 108 | 2 | -4457 | 0.03 | 0.000 |
| M7 Normative Q-value | 2 | 9992 | 1144 | 3 | -4962 | 0.09 | 0.002 |
| M8 Hybrid | 6 | 9110 | 261 | 0 | -4456 | 0.03 | 0.000 |

**Supplementary Table S2. Whole-brain clusters for GLMs 1–5.** All clusters surviving cluster-level FWE correction ($p < 0.05$; cluster-forming $p < 0.001$; sign-flip permutation test with 5,000 permutations). Peak coordinates in MNI space (mm); region, Neuromorphometrics label at or nearest to the peak; k, cluster size in 3-mm voxels. N = 25.

| Region | x | y | z | Peak t | k | p (FWE) |
|---|---|---|---|---|---|---|
| **Prospective value of the chosen card (positive)** | | | | | | |
| left precuneus | -12 | -67 | 38 | 5.63 | 1550 | < 0.001 |
| right cerebellum | 27 | -55 | -19 | 5.79 | 1239 | < 0.001 |
| left angular gyrus | -39 | -52 | 44 | 5.52 | 402 | 0.001 |
| left inferior occipital gyrus | -33 | -91 | -1 | 4.97 | 363 | 0.002 |
| left precentral gyrus | -27 | -7 | 59 | 5.08 | 242 | 0.002 |
| left lateral orbital gyrus | -33 | 50 | -13 | 4.84 | 132 | 0.006 |
| right anterior orbital gyrus | 24 | 53 | -16 | 4.20 | 118 | 0.007 |
| right basal forebrain / ventral striatum | 12 | 5 | -13 | 5.16 | 74 | 0.01 |
| left thalamus | -12 | -19 | 5 | 4.46 | 71 | 0.02 |
| right superior frontal gyrus | 27 | -4 | 62 | 4.83 | 66 | 0.02 |
| left accumbens area | -12 | 5 | -10 | 6.29 | 51 | 0.02 |
| left middle frontal gyrus | -45 | 14 | 44 | 4.98 | 41 | 0.03 |
| right thalamus | 9 | -7 | 2 | 4.62 | 35 | 0.04 |
| left angular gyrus | -54 | -55 | 23 | 4.35 | 32 | 0.04 |
| left supplementary motor cortex | -3 | -13 | 47 | 4.04 | 27 | 0.05 |
| **Immediate value of the chosen card (negative)** | | | | | | |
| left anterior insula | -30 | 20 | -10 | -4.46 | 75 | 0.009 |
| left supplementary motor cortex | 0 | 23 | 50 | -5.48 | 67 | 0.01 |
| right cuneus | 9 | -94 | 17 | -4.99 | 41 | 0.03 |
| **Immediate value of the chosen card (positive)** | | | | | | |
| right lingual gyrus | 3 | -85 | -7 | 11.34 | 860 | < 0.001 |
| left superior occipital gyrus | -18 | -85 | 32 | 4.67 | 162 | 0.003 |
| **Utility of the unchosen card (negative)** | | | | | | |
| right occipital pole | 12 | -94 | 23 | -6.23 | 406 | < 0.001 |
| right posterior insula | 45 | -7 | -1 | -4.98 | 265 | 0.002 |
| **Utility of the unchosen card (positive)** | | | | | | |
| right anterior insula | 33 | 20 | 2 | 5.03 | 84 | 0.02 |
| right superior frontal gyrus medial segment | 3 | 32 | 44 | 4.46 | 71 | 0.02 |
| right middle frontal gyrus | 42 | 23 | 50 | 4.30 | 36 | 0.04 |
| **Chosen minus unchosen utility (positive)** | | | | | | |
| right superior occipital gyrus | 21 | -88 | 32 | 6.53 | 2699 | < 0.001 |
| cerebellar vermal lobules viii-x | -6 | -70 | -43 | 4.93 | 272 | 0.001 |
| right superior temporal gyrus | 60 | 5 | -10 | 4.52 | 186 | 0.002 |
| left putamen | -21 | 5 | 8 | 5.09 | 126 | 0.006 |
| left middle cingulate gyrus | 0 | -22 | 29 | 5.29 | 121 | 0.007 |
| left caudate | -9 | 23 | 5 | 5.49 | 42 | 0.04 |
| right postcentral gyrus | 45 | -22 | 38 | 4.29 | 41 | 0.04 |
| **Prospective minus immediate value (negative)** | | | | | | |
| right lingual gyrus | 3 | -85 | -4 | -7.76 | 118 | 0.01 |
| **Prospective minus immediate value (positive)** | | | | | | |
| right superior parietal lobule | 30 | -64 | 35 | 5.42 | 557 | < 0.001 |
| left angular gyrus | -42 | -58 | 53 | 5.24 | 300 | 0.001 |
| right precentral gyrus | 48 | 8 | 32 | 4.97 | 188 | 0.003 |
| left middle frontal gyrus | -33 | 59 | -1 | 5.16 | 180 | 0.003 |
| left inferior occipital gyrus | -51 | -73 | -1 | 5.54 | 142 | 0.005 |
| right cuneus | 9 | -91 | 17 | 5.63 | 129 | 0.006 |
| left cerebellum | -39 | -49 | -31 | 4.91 | 121 | 0.007 |

| | | | | | | |
|---|---|---|---|---|---|---|
| right inferior temporal gyrus | 42 | -58 | -10 | 4.92 | 102 | 0.009 |
| left middle frontal gyrus | -45 | 14 | 44 | 4.87 | 97 | 0.01 |
| right lateral orbital gyrus | 36 | 59 | -7 | 4.27 | 76 | 0.02 |
| right anterior insula | 33 | 23 | -4 | 4.70 | 64 | 0.02 |
| left putamen | -27 | 14 | 2 | 5.31 | 61 | 0.02 |
| right posterior cingulate gyrus | 3 | -31 | 41 | 4.30 | 50 | 0.03 |
| right cerebellum | 15 | -64 | -19 | 4.27 | 41 | 0.04 |
| right superior occipital gyrus | 33 | -88 | 17 | 4.87 | 39 | 0.04 |
| **Signed decision value (chosen - unchosen) (negative)** | | | | | | |
| right anterior insula | 30 | 20 | -10 | -5.45 | 157 | 0.001 |
| left anterior insula | -30 | 26 | -4 | -4.78 | 105 | 0.003 |
| left supplementary motor cortex | 0 | 23 | 50 | -4.35 | 71 | 0.009 |
| right middle frontal gyrus | 42 | 20 | 50 | -4.62 | 50 | 0.02 |
| **Signed decision value (chosen - unchosen) (positive)** | | | | | | |
| right planum polare | 51 | 2 | -1 | 4.90 | 188 | 0.001 |
| left putamen | -24 | 8 | -1 | 4.74 | 53 | 0.01 |
| right cerebellum | 27 | -55 | -49 | 4.67 | 51 | 0.02 |
| left caudate | -9 | 23 | 2 | 4.79 | 43 | 0.02 |
| right occipital pole | 18 | -97 | 5 | 4.05 | 41 | 0.02 |
| right lingual gyrus | 6 | -91 | -7 | 4.03 | 37 | 0.03 |
| **Unsigned decision value (positive)** | | | | | | |
| left middle temporal gyrus | -54 | -64 | 11 | 4.76 | 82 | 0.01 |
| right anterior insula | 30 | 20 | -10 | 4.83 | 62 | 0.02 |
| right superior frontal gyrus medial segment | 3 | 41 | 32 | 4.54 | 51 | 0.02 |
| right lingual gyrus | 24 | -52 | 2 | 4.00 | 45 | 0.03 |
| right middle frontal gyrus | 42 | 11 | 44 | 4.28 | 41 | 0.03 |
| left anterior insula | -30 | 20 | -7 | 4.56 | 32 | 0.04 |
| **Difficulty (-unsigned decision value) (negative)** | | | | | | |
| left middle temporal gyrus | -54 | -64 | 11 | -4.76 | 82 | 0.01 |
| right anterior insula | 30 | 20 | -10 | -4.83 | 62 | 0.02 |
| right superior frontal gyrus medial segment | 3 | 41 | 32 | -4.54 | 51 | 0.02 |
| right lingual gyrus | 24 | -52 | 2 | -4.00 | 45 | 0.03 |
| right middle frontal gyrus | 42 | 11 | 44 | -4.28 | 41 | 0.03 |
| left anterior insula | -30 | 20 | -7 | -4.56 | 32 | 0.04 |
| brain stem | 3 | -16 | -19 | -4.62 | 30 | 0.04 |
| **Conflict trials: lower vs higher card chosen (positive)** | | | | | | |
| left supramarginal gyrus | -54 | -52 | 47 | 4.89 | 47 | 0.02 |
| **Lower card chosen: conflict vs neutral (positive)** | | | | | | |
| left postcentral gyrus | -42 | -22 | 62 | 4.83 | 52 | 0.01 |
| **Lower choice x conflict (positive)** | | | | | | |
| left postcentral gyrus | -39 | -25 | 65 | 5.36 | 96 | 0.007 |
| right cerebellum | 24 | -55 | -19 | 4.38 | 68 | 0.01 |
| **Prospective value (goal gradient) (positive)** | | | | | | |
| left precuneus | -12 | -67 | 38 | 5.75 | 1452 | < 0.001 |
| left cerebellum | -39 | -49 | -31 | 5.96 | 1188 | < 0.001 |
| left angular gyrus | -42 | -61 | 53 | 5.83 | 582 | < 0.001 |
| left inferior occipital gyrus | -33 | -91 | -1 | 5.01 | 330 | 0.001 |
| right basal forebrain / ventral striatum | 12 | 5 | -13 | 5.08 | 185 | 0.004 |
| right inferior occipital gyrus | 33 | -91 | -1 | 4.93 | 182 | 0.004 |
| left precentral gyrus | -27 | -7 | 59 | 5.21 | 171 | 0.004 |
| left lateral orbital gyrus | -36 | 53 | -13 | 5.21 | 144 | 0.006 |
| right superior frontal gyrus | 27 | -4 | 59 | 5.14 | 126 | 0.007 |
| right anterior orbital gyrus | 27 | 59 | -10 | 4.21 | 122 | 0.007 |
| left middle frontal gyrus | -45 | 14 | 44 | 5.73 | 94 | 0.009 |

| | | | | | | |
|---|---|---|---|---|---|---|
| left accumbens area | -12 | 5 | -10 | 6.03 | 49 | 0.03 |
| right anterior insula | 33 | 17 | -7 | 5.33 | 41 | 0.03 |
| left middle temporal gyrus | -63 | -40 | -7 | 4.34 | 36 | 0.04 |
| **Immediate value (goal gradient) (negative)** | | | | | | |
| right cuneus | 9 | -94 | 17 | -5.49 | 114 | 0.004 |
| left supplementary motor cortex | 0 | 23 | 50 | -5.17 | 32 | 0.05 |
| **Immediate value (goal gradient) (positive)** | | | | | | |
| right lingual gyrus | 3 | -85 | -7 | 11.28 | 669 | < 0.001 |
| **Expected set bonus (planner) (positive)** | | | | | | |
| left occipital fusiform gyrus | -18 | -91 | -7 | 8.02 | 4591 | < 0.001 |
| left precentral gyrus | -30 | -7 | 56 | 5.59 | 565 | < 0.001 |
| right middle frontal gyrus | 27 | 2 | 59 | 5.37 | 209 | 0.003 |
| cerebellar vermal lobules viii-x | -3 | -55 | -34 | 5.25 | 137 | 0.007 |
| left thalamus | -6 | 2 | 5 | 5.34 | 94 | 0.01 |
| right precentral gyrus | 45 | 5 | 32 | 5.38 | 76 | 0.02 |
| right middle frontal gyrus | 48 | 35 | 29 | 4.68 | 62 | 0.02 |
| right frontal pole | 30 | 65 | 8 | 4.67 | 52 | 0.03 |
| right medial orbital gyrus | 18 | 47 | -22 | 4.42 | 43 | 0.03 |
| left thalamus | 0 | -19 | 2 | 4.57 | 43 | 0.03 |
| right anterior insula | 33 | 23 | -4 | 5.21 | 39 | 0.04 |
| left middle frontal gyrus | -27 | 62 | -4 | 3.86 | 38 | 0.04 |
| **Points (planner) (negative)** | | | | | | |
| right cuneus | 9 | -94 | 17 | -5.50 | 117 | 0.004 |
| **Points (planner) (positive)** | | | | | | |
| right lingual gyrus | 3 | -85 | -7 | 11.28 | 647 | < 0.001 |

**Supplementary Table S3. Cross-validated Bayesian comparison of the goal-gradient (GLM 4) and planner (GLM 5) GLMs.** Share, participants with higher cross-validated log model evidence for GLM 4; RFX frequency, expected frequency of GLM 4 from random-effects Bayesian model selection; summed log BF, sum over participants of the log Bayes factor for GLM 4 versus GLM 5 (positive values favor the goal-gradient GLM). N = 25.

| ROI | Share favoring GLM 4 | RFX frequency | Summed log BF |
|---|---|---|---|
| vmPFC | 60% | 0.67 | +34.6 |
| striatum | 68% | 0.73 | +37.9 |
| L caudate | 72% | 0.75 | +33.6 |
| R caudate | 68% | 0.67 | +35.8 |
| L insula | 56% | 0.66 | +49.6 |
| R insula | 60% | 0.62 | +40.3 |
| precuneus | 56% | 0.69 | +38.6 |

**Supplementary Table S4. Temporal signal-to-noise ratio in the regions of interest.** Mean, standard deviation and minimum across participants of each participant's mean tSNR in the ROI; coverage, the smallest share of ROI voxels inside a participant's analysis mask. N = 25.

| ROI | Mean tSNR | SD | Minimum | Coverage |
|---|---|---|---|---|
| vmPFC | 49 | 6 | 38.3 | 100% |
| striatum | 38.9 | 3.4 | 32.5 | 100% |
| L caudate | 44.8 | 4.3 | 36.3 | 100% |
| R caudate | 46.7 | 3.8 | 38.8 | 100% |
| L insula | 49.9 | 4.8 | 42.3 | 100% |
| R insula | 47.7 | 3.3 | 39.7 | 100% |
| precuneus | 71.4 | 7.7 | 57.8 | 100% |

**Supplementary Table S5. Mixed-effects logistic regressions of choice.** Upper part: model comparison of regressions of the choice of the right card on the right-minus-left differences in points (valuedif), feasible suit sets (suitdif), feasible digit sets (digitdif) and all feasible sets (setdif), each with participant random slopes. Lower part: fixed effects of the best model (b5). N = 29 participants, 8,525 trials.

| Model | Fixed effects | BIC | ΔBIC |
|---|---|---|---|
| b1 | valuedif | 9558.5 | 475.7 |
| b2 | suitdif + digitdif | 10932.4 | 1849.7 |
| b3 | valuedif + suitdif | 9262.4 | 179.6 |
| b4 | valuedif + digitdif | 9444.9 | 362.1 |
| b5 | valuedif + suitdif + digitdif | 9082.8 | 0.0 |
| b6 | valuedif + setdif | 9100.3 | 17.5 |

| Term (model b5) | Estimate | SE | z | p |
|---|---|---|---|---|
| (Intercept) | -0.043 | 0.040 | -1.07 | 0.28 |
| valuedif | 0.600 | 0.077 | 7.75 | < 0.001 |
| suitdif | 1.055 | 0.128 | 8.21 | < 0.001 |
| digitdif | 0.701 | 0.094 | 7.44 | < 0.001 |

**Supplementary Table S6. Model-based GLMs.** All GLMs modeled choices at stimulus onset with duration equal to the response time. Model-based GLMs (1, 2, 4, 5) also contained four control modulators (trial within the game, trial within the session, response time, response button); every GLM contained six realignment parameters per run. Utilities are from the goal-gradient model with group-median value and set weights and each participant's own gradient steepness κ. In the analysis code, GLMs 1–5 are named GLM1_components, GLM2_decision_value, GLM3_conflict, GLM4_goalgradient and GLM5_planner.

| **GLM** | **Regressors of interest** | **Contrasts** |
|---|---|---|
| 1 Value components | Immediate utility of the chosen card; prospective utility of the chosen card; utility of the unchosen card | Each regressor; prospective − immediate; chosen − unchosen utility |
| 2 Decision value | Signed decision value (chosen − unchosen utility); unsigned decision value | Each regressor |
| 3 Conflict | Conditions: congruent, neutral and conflict trials, each split by higher- or lower-point choice; ties | Conflict lower vs higher; lower: conflict vs neutral; interaction; conflict vs congruent |
| 4 Goal gradient | Immediate utility; goal-gradient prospective utility | Each regressor; MACS vs GLM 5 |
| 5 Planner | Points of the chosen card; expected set bonus (planner) | Each regressor; MACS vs GLM 4 |
